# Comparative Study of Photodynamic and Sonodynamic Therapy Using a PSMA-Targeted Up-conversion Nanoplatform for Castration-Resistant Prostate Cancer

**DOI:** 10.64898/2026.04.10.717693

**Authors:** Mehrgan Ghazaeian, Rahul Kumar Das, Ravindra K. Pandey, Supriya D. Mahajan, Mykhaylo Dukh, Yanda Cheng, Andrey Kuzmin, Artem Pliss, Stanley A. Schwartz, Jun Xia, Ravikumar Aalinkeel, Paras N. Prasad

## Abstract

Photodynamic therapy (PDT) for prostate cancer is limited by the shallow penetration of visible light into deep-seated tumors. In contrast, sonodynamic therapy (SDT) enables noninvasive deep-tissue activation using ultrasound, while near-infrared photodynamic therapy (NIR-PDT) enhances tissue penetration through up-conversion-mediated activation. Based on these advantages, we developed a prostate-specific membrane antigen (PSMA)-targeted nanotheranostic platform (UCNPs@mSiO₂/HPPH@TCS) to compare NIR-PDT and SDT in vitro for penetration-enhanced and targeted prostate cancer therapy. The nanoformulation consists of NaYF_4_:Yb^3+^, Er^3+^ up-conversion nanoparticles coated with mesoporous silica for high loading of HPPH (Photochlor), which functions as both photosensitizer and sonosensitizer, and a PSMA-targeted chitosan shell for selective tumor targeting. Physicochemical characterization confirms a uniform core-shell structure (∼63 nm). Tissue-mimicking depth studies demonstrate that SDT and NIR-PDT achieve greater penetration than conventional 665 nm PDT in thicker tissue layers. Intracellular reactive oxygen species (ROS) analysis shows that SDT generates higher total ROS levels, whereas NIR-PDT produces greater singlet oxygen generation. Three-dimensional spheroid models further validate efficacy, demonstrating rapid spheroid collapse via apoptosis, with SDT inducing earlier apoptosis and more uniform penetration than PDT. Collectively, the results suggest SDT provides superior overall therapeutic efficacy compared with NIR-PDT, highlighting the potential of this nanoplatform for precision prostate cancer therapy.

## 1. Introduction

Cancer remains a major global public health challenge and is currently among the leading causes of mortality worldwide.^[1, 2]^ Globally, prostate cancer (PCa) is the second most diagnosed malignancy in men after lung cancer, with particularly high incidence rates in South America, Western and Northern Europe, Australia, and North America.^[2]^ Standard therapeutic approaches for PCa include radical prostatectomy, radiotherapy, androgen-deprivation therapy, chemotherapy, and, in selected cases, immunotherapy, and are determined by disease stage and clinical context.^[3]^ However, these modalities often lack tumor specificity, resulting in unintended damage to surrounding healthy tissues and significant treatment-related side effects.^[4]^ Consequently, there remains a pressing need for selective, minimally invasive therapeutic strategies capable of improving therapeutic efficacy while minimizing systemic toxicity in PCa.

Among emerging alternatives, photodynamic therapy (PDT) and sonodynamic therapy (SDT) have gained increasing attention as non-invasive cancer treatment modalities.^[5]^ PDT induces selective tumor cell death through the generation of reactive oxygen species (ROS) following the activation of a photosensitizer (PS) in the presence of light and molecular oxygen.^[1, 6, 7]^ The resulting singlet oxygen (¹O₂) and other ROS cause oxidative damage to cellular membranes, proteins, and nucleic acids, ultimately leading to apoptosis. Among second-generation PSs, 2-[1-hexyloxyethyl]-2-devinyl pyropheophorbide (HPPH) exhibits favorable photophysical properties, strong absorption within the therapeutic window, enhanced tumor selectivity, and reduced cutaneous photosensitivity compared to earlier PSs.^[8]^ Despite these advantages, the clinical translation of PDT remains constrained by limited light penetration depth and suboptimal PS solubility, stability, and pharmacokinetics, particularly in deep-seated tumors.^[1, 9]^

To overcome the penetration limitation of conventional PDT, SDT has emerged as a complementary non-invasive approach. SDT employs low-intensity ultrasound (US) to activate sonosensitizers, enabling ROS generation in deep tissues with minimal attenuation.^[10–13]^ Upon US exposure, acoustic cavitation and associated mechanical and chemical effects trigger oxidative stress, leading to cancer cell damage and death. A broad range of sonosensitizers, including tetrapyrrole-based compounds like HPPH, small-molecule drugs, and nanomaterials, have been explored, with US-responsive systems offering significant advantages for treating tumors inaccessible to light-based therapies.^[11, 14]^

In parallel with SDT, up-conversion nanoparticle (UCNPs)-mediated PDT has emerged as an alternative strategy to extend photodynamic activation into deeper tissues. UCNPs function as nanoscale optical transducers, converting deeply penetrating near-infrared (NIR) light into higher-energy visible emissions capable of activating conventional PSs.^[15]^ Because NIR light lies within the biological optical transparency window, UCNPs-mediated PDT enables deeper tissue penetration while minimizing photodamage to surrounding normal tissues, which do not absorb at NIR^. [16, 17]^ Importantly, this strategy allows PDT to partially overcome its intrinsic depth limitation and provides a direct framework for comparing NIR-driven PDT with SDT, both of which aim to achieve effective ROS generation in deep tumor regions through distinct physical activation mechanisms.

Despite these advances, conventional PSs, being generally hydrophobic, still suffer from poor aqueous solubility, low tumor accumulation, and limited in vivo stability. Nanotechnology-based delivery platforms address these challenges by improving aqueous dispersibility, pharmacokinetics, tumor selectivity, and therapeutic versatility.^[1]^ Rare-earth lanthanide-doped UCNPs are particularly attractive for biomedical applications due to their photostability, sharp emission bands, resistance to photobleaching, and ability to produce NIR-to-visible up-conversion.^[18]^ For enhanced biocompatibility and drug-loading capacity, UCNPs are commonly coated with mesoporous silica (mSiO₂), which provides tunable pore structures and surface functionalization with –OH, –COOH, –NH₂, or –PO₄ groups, enabling high payload retention and controlled release.^[19, 20]^ Furthermore, chitosan-based coatings offer additional advantages, including pH-responsive drug release, improved biocompatibility, and the potential for tumor-specific targeting.^[21, 22]^ In PCa, active targeting can be introduced through prostate-specific membrane antigen (PSMA) ligands, particularly glutamate-urea derivatives, which exhibit high binding affinity toward PSMA-overexpressing PCa cells.^[23–25]^

Three-dimensional (3D) tumor spheroid models have emerged as an essential intermediate platform between conventional monolayer cultures and in vivo tumor systems for therapy evaluation, as they more accurately recapitulate the architectural, metabolic, and diffusion constraints of solid tumors.^[26]^ Moreover, the dense extracellular matrix and cell-cell interactions within spheroids impose significant barriers to nanoparticle penetration and distribution, thereby providing a more stringent and clinically relevant assessment of delivery efficiency and therapeutic response.^[27]^ Given that both PDT and SDT rely on the spatiotemporal generation of ROS within tumor tissues, evaluation in 3D spheroid systems is necessary to capture treatment resistance mechanisms arising from hypoxia and limited nanoparticle diffusion, which are not adequately modeled in 2D systems. Therefore, assessing PDT and SDT efficacy in spheroid assays is critical for accurately evaluating the therapeutic potential and translational relevance of advanced nanoplatforms designed for combined PDT/SDT applications.

Based on these principles, we report the design of a multifunctional PSMA-targeted UCNPs@mSiO₂/HPPH nanoplatform that integrates deep-tissue NIR activation, high photosensitizer loading, and tumor-selective targeting (Figure 1a). By integrating up-conversion-mediated PDT and US mediated SDT within a single nanocomposite architecture utilizing a single sensitizer, this platform facilitates a direct, penetration-matched comparative analysis of light- and ultrasound-activated therapeutic modalities. Our findings demonstrate that this PSMA-targeted nanoformulation (NF) enables robust therapeutic responses in PCa cells, with SDT producing superior intracellular ROS generation and enhanced efficacy relative to UCNP-mediated PDT, while maintaining minimal photothermal interference. Crucially, we provide the first systematic comparison of PDT and SDT within a 3D in vitro spheroid architecture, demonstrating that SDT achieves more uniform spheroid penetration and accelerates structural degradation compared to PDT. Collectively, these results underscore the importance of benchmarking complementary activation strategies to overcome the intrinsic limitations of conventional PDT, establishing a new framework for precision nanotherapeutics in deep-seated prostate cancer treatment.

**Figure 1.**
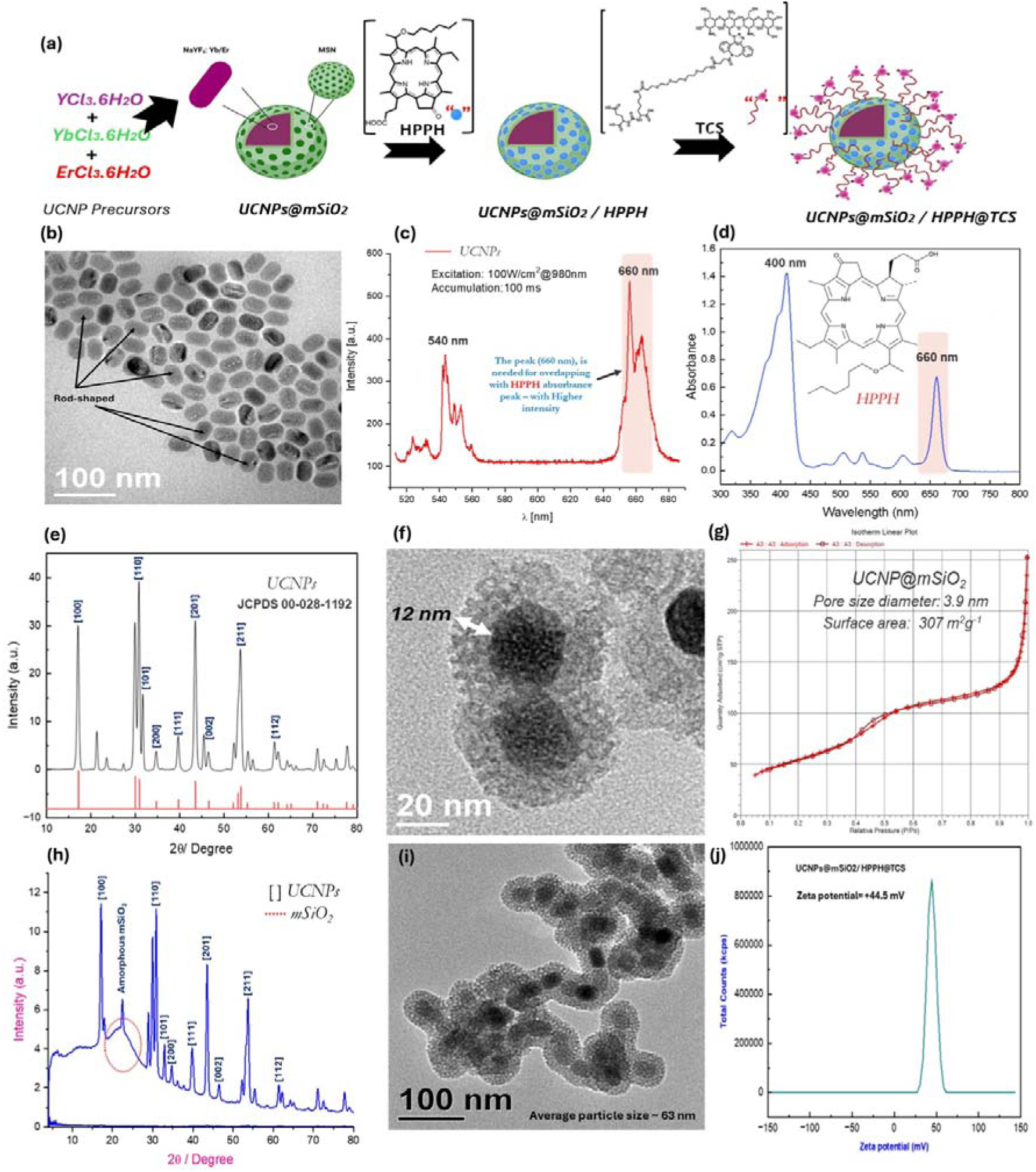
Synthesis and Physico-chemical Characterization of UCNPs@mSiO2/HPPH@TCS. (a) Schematic illustration of the stepwise synthesis and surface functionalization of the PSMA-targeted UCNPs@mSiO₂ nanoformulation. UCNP cores were first prepared via thermal decomposition, followed by silica coating using CTAB as a template. Mesopores were generated after template removal. The photosensitizer HPPH was loaded into the mesopores, and the nanoparticle surface was functionalized with PSMA-targeted chitosan to enable active targeting. (b) TEM image of NaYF_4_: Yb^3+^, Er^3+^ UCNPs, displaying uniform rod-like morphology (average size: 40 ± 3 nm). (c) Photoluminescence emission spectrum of UCNPs under 980 nm excitation, showing dominant peaks at 540 and 660 nm; the latter spectrally overlaps with the (d) characteristic Q-band absorbance of HPPH centered at ∼665 nm. (e) X-ray diffraction (XRD) pattern of NaYF4: Yb, Er UCNPs, confirming the high crystallinity and phase purity of the hexagonal (β-phase) structure. All diffraction peaks are in excellent agreement with the standard hexagonal NaYF_4_ reference pattern (JCPDS No. 00-028-1192), with no detectable peaks from the cubic (α-phase) or other impurities. (f) TEM micrograph of UCNPs@mSiO_2_ core-shell nanoparticles, illustrating a uniform ∼12 nm silica shell and a total diameter of ∼52 nm. (g) Nitrogen adsorption–desorption isotherms and pore size distribution of UCNPs@mSiO_2_, indicating a high specific surface area (307 m^2^g^-1^) and uniform mesopores (∼3.9 nm). (h) XRD pattern of UCNPs@mSiO_2_ with a broad diffraction peak at 2ө ∼ 22°, confirming the amorphous nature of the silica coating. (i) TEM image of the final PSMA-targeted chitosan-capped nanoparticles (total diameter: ∼63 nm). (j) Zeta potential distribution of the final nanoformulation, showing a stable surface charge of +44.5 mV.

## 2. Results and Discussion

### 2.1. Characterizations

#### 2.1.1. Characterizations of UCNPs

To establish the structural, morphological, and optical attributes required for efficient up-conversion-mediated PDT, the synthesized UCNPs were systematically characterized using transmission electron microscopy (TEM) and high-resolution TEM (HRTEM), powder X-ray diffraction (XRD), and photoluminescence (PL) spectroscopy. TEM revealed that the synthesized UCNPs were predominantly rod-shaped with a uniform size distribution of approximately 40 ± 3 nm (Figure 1b, Figure S1). Anisotropic core-shell nanostructures, particularly rod-shaped UCNPs, offer distinct advantages as FRET donors because they minimize the distance between internal activators and surface-bound photosensitizers. As recently reported, these rod-shaped morphologies exhibit more robust up-conversion emissions and reduced surface quenching compared to isotropic spheres or disks, leading to accelerated structural degradation of tumor models during treatment.^[28]^ PL spectra under 975 nm excitation exhibited characteristic Er³⁺ emissions with strong green (∼540 nm; ²H₁₁/₂ → ⁴I₁₅/₂) and red (∼658 nm; ⁴F₉/₂ → ⁴I₁₅/₂) peaks (Figure 1c). The enhanced 658 nm red emission observed with higher Yb³⁺ doping (40%) resulted from more efficient Yb³⁺→Er³⁺ energy transfer and aligns well with the absorption band of HPPH (Figure 1d), thereby improving excitation efficiency for PDT. Collectively, the structural and optical analyses confirm that the synthesized β-NaYF₄: Yb³⁺, Er³⁺ UCNPs possess optimal morphology, crystallinity, and luminescent properties for use in PDT applications.

The crystalline phase and structural purity of the UCNPs were confirmed by XRD analysis, which showed diffraction peaks matching the standard JCPDS card No. 00-028-1192, corresponding to hexagonal β-NaYF₄ (Figure 1e). The sharp peaks indicated high crystallinity, and only minor additional peaks were observed, likely due to trace impurities or residual α-phase, without affecting the overall dominance of the β-phase. The incorporation of Yb³⁺ and Er³⁺ dopants maintained the hexagonal symmetry, confirming their successful substitution at Y³⁺ sites.

#### 2.1.2. Characterization of UCNPs@mSiO₂ Nanoparticles

TEM analysis further confirmed the formation of a uniform core–shell structure, with a silica shell approximately 12 nm thick surrounding the rod-shaped UCNP cores, yielding an average total particle size of about 52 nm (Figure 1f). Fourier Transform Infrared (FTIR) spectra (Figure S2 (a, b)) of UCNPs@mSiO₂ exhibited characteristic bands at 1070 cm⁻¹ and 800 cm⁻¹, corresponding to Si–O–Si asymmetric and symmetric stretching vibrations, respectively, confirming the formation of a silica network on the nanoparticle surface. Peaks at 2850 cm⁻¹ and 2920 cm⁻¹, attributed to the C–H stretching vibrations of the Cetyltrimethylammonium bromide (CTAB) surfactant, disappeared after template removal, indicating successful elimination of CTAB and the formation of clean mesoporous channels. The coating was uniform and well-defined, suggesting efficient silica growth and preservation of particle morphology. Dynamic Light Scattering (DLS) measurements revealed a hydrodynamic diameter of 197.6 nm and a zeta potential of –13.68 mV, consistent with the presence of negatively charged silanol (–SiOH) groups on the silica surface, which enhance colloidal stability and enable further surface functionalization (Figure S3). These results collectively confirm the successful synthesis of uniform, stable UCNPs@mSiO₂ nanoparticles with preserved crystallinity and well-formed mesoporous silica shells suitable for subsequent functionalization and therapeutic applications.

Nitrogen adsorption–desorption analysis was performed to evaluate the textural properties of the UCNPs after mesoporous silica coating. The resulting isotherm exhibited characteristic features of mesoporous materials, confirming the successful formation of a porous silica shell. The Brunauer–Emmett–Teller (BET) specific surface area was determined to be 307 m² g⁻¹, indicating a highly accessible surface suitable for efficient PS loading. Furthermore, Barrett–Joyner–Halenda (BJH) pore size analysis revealed a narrow pore size distribution with an average pore diameter of approximately 3.9 nm, which is well aligned with the molecular dimensions of HPPH and supports effective encapsulation while preserving mass transport and diffusion properties (Figure 1g).

XRD analysis confirmed that the core retained the hexagonal β-NaYF₄ crystal structure (JCPDS No. 00-028-1192) after silica coating, while an additional broad peak around 2θ ≈ 22° indicated the presence of amorphous silica (Figure 1h).

#### 2.1.3. Characterization of amine-modified UCNPs@mSiO_2_ (UCNPs@mSiO_2_-NH_2_)

To provide reactive handles for subsequent shell conjugation and drug loading, the silica surface of the UCNPs@mSiO_2_ was functionalized with primary amine groups using (3-Aminopropyl) triethoxysilane (APTES). The successful surface modification of UCNPs@mSiO₂ with amine groups was confirmed through FTIR, TEM, and DLS analyses. The FTIR spectrum of UCNPs@mSiO₂–NH₂ (Figure S4) retained the characteristic Si–O–Si stretching bands at 1070 and 800 cm⁻¹ from the silica network, while a new distinct peak appeared at 1630 cm⁻¹, corresponding to the N–H bending vibration of the amine groups (–NH₂). The emergence of this peak confirms the successful grafting of APTES molecules onto the silica surface. DLS measurements revealed an increase in the hydrodynamic diameter from 197.6 nm to 229 nm, accompanied by a shift in zeta potential from –13.68 mV to +32 mV (Figure S5). This significant change in surface charge reflects the introduction of positively charged amine groups, which enhance colloidal stability and provide reactive sites for subsequent conjugation steps. Collectively, these results confirm the effective amine functionalization of the UCNPs@mSiO₂ nanoparticles while maintaining their structural and colloidal stability.

#### 2.1.4. Drug Loading and Characterization of UCNPs@mSiO₂–NH₂/HPPH Nanoparticles

The loading of the PS (HPPH) onto amine-functionalized UCNPs@mSiO₂ nanoparticles was achieved through covalent coupling using (1-Ethyl-3(3-dimethylaminopropyl) carbodiimide and N-hydroxysuccinimide) (EDC/NHS) chemistry. UV/Vis spectrophotometry was used to quantify the loading efficiency. Based on the absorbance of unbound HPPH in the supernatant at 660 nm. The drug loading efficiency (DLE) was calculated using the equation below:

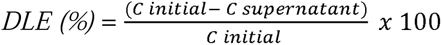

where *C* initial and *C* supernatant represent the initial and residual concentrations of HPPH, respectively. Based on this calculation, the loading efficiency was determined to be 92%, indicating highly effective covalent conjugation of HPPH onto the nanoparticle surface. This high efficiency highlights the strong affinity and stability of the EDC/NHS-mediated amide bond formation between the –NH₂ groups of APTES and the –COOH groups of HPPH. DLS analysis revealed an increase in hydrodynamic diameter from 229 nm (UCNPs@mSiO₂-NH₂) to 267 nm after HPPH loading, confirming successful surface conjugation and an increase in molecular weight (Figure S6 (a)). The zeta potential decreased from +32 mV to +16.1 mV, reflecting the partial neutralization of surface amine groups upon amide bond formation (Figure S6 (b)). These observations collectively confirm the efficient covalent attachment of HPPH to the UCNPs@mSiO₂-NH₂ nanoparticles, yielding a stable nanoconjugate with high drug loading capacity, an essential feature for enhanced ROS generation and improved PDT efficacy.

#### 2.1.5. Characterization of PSMA-targeted chitosan-capped UCNPs@mSiO_2_/HPPH

The characterization of the final nanocomposite, after coating UCNPs@mSiO₂/HPPH nanoparticles with PSMA-targeted chitosan previously synthesized in our laboratory, was performed to confirm successful functionalization. TEM analysis revealed that the NF exhibited an average particle size of approximately 63 nm following chitosan coating around the silica shell (Figure 1i). The silica shell also appeared denser and more uniform, indicating enhanced structural integrity imparted by the polymer coating. DLS and zeta potential analyses further confirmed the successful modification. The hydrodynamic size increased to 272 nm (Figure S7), while the zeta potential rose to +44.5 mV due to the positively charged amino groups of chitosan (Figure 1j). This high positive surface charge verifies successful coating and enhances colloidal stability through electrostatic repulsion. Overall, these results confirm the effective surface functionalization of UCNPs@mSiO₂/HPPH with PSMA-targeted chitosan, producing a stable and well-defined final nanocomposite suitable for targeted biomedical applications, which we define as UCNPs@mSiO₂/HPPH@TCS.

### 2.2. Acellular Studies

#### 2.2.1. Optimization of 980 nm Laser Parameters and Photothermal Safety Evaluation

Response-Surface Methodology (RSM) implemented in Design-Expert^®^ software was employed to model and describe the relationship between 980 nm laser irradiation parameters and temperature elevation. Based on the experimental design generated by the software, the photothermal performance of the synthesized nanoparticles was systematically evaluated in an acellular suspension under NIR laser irradiation using an IR thermal imaging camera. This approach enabled the identification of an optimal irradiation window that ensures efficient excitation of UCNPs for PDT, while minimizing unintended photothermal heating that could confound biological outcomes in subsequent in vitro studies.

##### Response-Surface Modeling and Statistical Validation

To quantitatively describe the relationship between laser parameters and temperature elevation, RSM was applied using Design-Expert software. Fit summary analysis for the temperature response identified a cubic polynomial model as the most appropriate. The selected model was statistically significant (p = 0.0003) with an insignificant lack of fitness, indicating good agreement between the model and experimental data. The high coefficient of determination (R² = 0.9985) further confirms the robustness and predictive accuracy of the model (Table S1).^[29]^ Model validity was assessed using a predicted versus actual temperature plot (Figure S8(a)), which showed excellent correlation, with data points closely distributed along the parity line. This strong agreement confirms that the model reliably captures the photothermal behavior of the nanoparticle system across the tested irradiation conditions.

##### Interaction Between Irradiation Time and Laser Intensity

The interaction plot illustrates the combined effects of irradiation time and laser intensity on temperature elevation, as shown in Figure S8(b). At the higher laser intensity (1.0 W= 1.27 W. cm^-2^), the temperature increased sharply with irradiation time, rapidly surpassing physiological thresholds. In contrast, at (0.5 W= 0.63 W.cm^-2^), temperature increases were modest and plateaued even at extended exposure times. The non-parallel interaction curves highlight a significant interaction between time and intensity, emphasizing that photothermal behavior is governed by their combined influence rather than either factor alone.

##### Numerical Optimization and Selection of PDT Conditions

Numerical optimization was performed to maintain the maximum temperature below 37 °C while permitting sufficient irradiation duration for effective UCNPs excitation. Under this constraint, Design-Expert identified an optimal solution with a desirability value of 1.00. The predicted optimal parameters were an irradiation time of 162.123 seconds and a laser intensity of 0.5 W, corresponding to a temperature of 31.24 °C, well within the defined safety window.^[30]^ Based on these findings, a laser intensity of 0.5 W (0.63 W.cm^-2^) was selected for all subsequent cellular PDT experiments, with a maximum irradiation time of 180 s. These conditions minimize photothermal effects and ensure that any observed cytotoxicity arises predominantly from photodynamically generated reactive oxygen species rather than from nonspecific thermal damage (Figure S9).

#### 2.2.2. Optimization of SDT Parameters Using Response-Surface Methodology

The Determinant-optimal (D-optimal) response surface experiment identified nonlinear relationships and interactions between US intensity and time that determine ROS generation from our NF. Applying a base 10 log transformation improved adherence to ANOVA assumptions. It produced a statistically significant quadratic model (p = 0.0316) with a high R^2^ (0.9379) and reasonable adjusted R^2^ (0.7764), indicating the model explains a large fraction of the transformed response variance while accounting for the number of terms in the model (Table S2).^[31]^ The predicted vs actual plot showed that most points lie near the 45° line, indicating good agreement between measured ROS values and model predictions; a few deviations at the highest and lowest responses suggest regions where experimental variability or minor model misspecification exist (Figure S10(a)). The presence of replicate and lack of fit points strengthened the ability to separate pure experimental error from lack of fit; the model passed the lack of fit checks reported by the software, supporting its adequacy over the tested region. The 3D surface visualizations reveal a response surface with pronounced curvature: ROS generation tends to increase at extreme combinations of factors rather than being strictly linear with intensity or time. High intensity (5 W.cm^-2^) combined with relatively short to moderate irradiation time produced high predicted ROS, whereas intermediate combinations often produced lower ROS (a central “valley” in the surface). Frequency acted as a categorical shift in the model, and optimization indicated that 3 MHz gave higher ROS under the tested conditions than 1 MHz. These patterns are consistent with a quadratic model in which both main effects and their interaction contribute substantially to the response (Figure S10 (b)). Mechanistically, such nonlinear dependence can arise because ultrasonic cavitation, mechanical effects, and sonosensitizer excitation depend on both acoustic intensity and exposure duration in a non-additive manner: short, intense pulses may favor efficient production of short-lived ROS, while extended exposure at intermediate intensity may increase scavenging or induce degradation of the sensitizer, reducing net ROS.^[32]^

##### Numerical optimization and selected operating point

Numerical optimization was configured to maximize ROS (targeting an upper bound of 100% in the software). The software returned to a single optimal solution with desirability = 1.000: frequency = 3 MHz, intensity = 5 W.cm^-2^, and time = 2 min. A desirability of 1 indicates that, within the model and the defined goals/constraints, the optimizer found a point that fully satisfies the target criteria. This optimal point sits at the high intensity edge with a relatively short exposure time, consistent with the acellular response surface that favors high intensity but non-prolonged irradiation (Figure S11).

##### Cellular validation and biological interpretation

To translate the acellular optimization into a cellular context, we tested acoustic conditions centered on the software’s recommendation (frequency fixed at 3 MHz) across intensities and times. In PC-3 cells, SDT with the NF did not result in a statistically significant reduction in viability across the tested conditions, reflecting the absence of PSMA-mediated nanoparticle uptake and, consequently, reduced intracellular therapeutic activation rather than inherent cellular resistance. In contrast, PSMA^+^ LNCaP cells displayed pronounced treatment sensitivity, consistent with efficient targeted uptake and enhanced intracellular therapeutic activation. The greatest cell killing occurred at intensity = 5 W.cm^-2^ and time = 3 min, a condition very close to the acellular optimum (5 W.cm^-2^, 2 min). This concordance between acellular prediction and cellular response in LNCaP supports the utility of the 1,3-diphenylisobenzofuran (DPBF) assay and the response surface model for identifying biologically relevant sonication settings. The difference in response between PC-3 and LNCaP cells, apart from targeted uptake, can also reflect several biological factors, intracellular localization, differences in antioxidant defenses (e.g., glutathione levels, catalase activity), differential membrane properties affecting cavitation-mediated effects, or distinct cell death pathways activated by ROS.^[33]^ The slightly longer optimal time observed in LNCaP (3 min vs 2 min predicted acellular) is reasonable given that intracellular diffusion, scavenging, and the physical environment within cells can alter effective ROS exposure and kinetics relative to an acellular solution.

#### 2.2.3. Infrared Thermal Imaging Analysis of Nanoparticle Photothermal Response under 980 nm NIR Irradiation (Figure 2a)

##### High-Power Screening (1.0 W (1.27 W.cm^-2^))

Initial screening was performed at a laser power of 1.0 W (1.27 W.cm^-2^) to assess the upper thermal response of the NP suspension (Figure 2b). Temperature increased markedly with irradiation time, rising from 29.0 °C at 15 s to 35.6 °C at 30 s and 36.8 °C at 60 s. Prolonged exposure resulted in a rapid transition into a hyperthermic regime, with temperatures exceeding physiological limits after 120 s (42.8 °C) and reaching a maximum of 47.5 °C at 180 s. These results demonstrate that, at 1.0 W, the nanoparticles exhibit a pronounced photothermal conversion capability. While such heating may be advantageous for photothermal therapy, it is unsuitable for isolating photodynamic effects, as temperatures above ∼42 °C can induce direct thermal cytotoxicity.^[34]^ Under these conditions, thermally driven cell death would dominate and obscure ROS-mediated photodynamic mechanisms, rendering this power setting inappropriate for PDT studies.^[34]^

**Figure 2.**
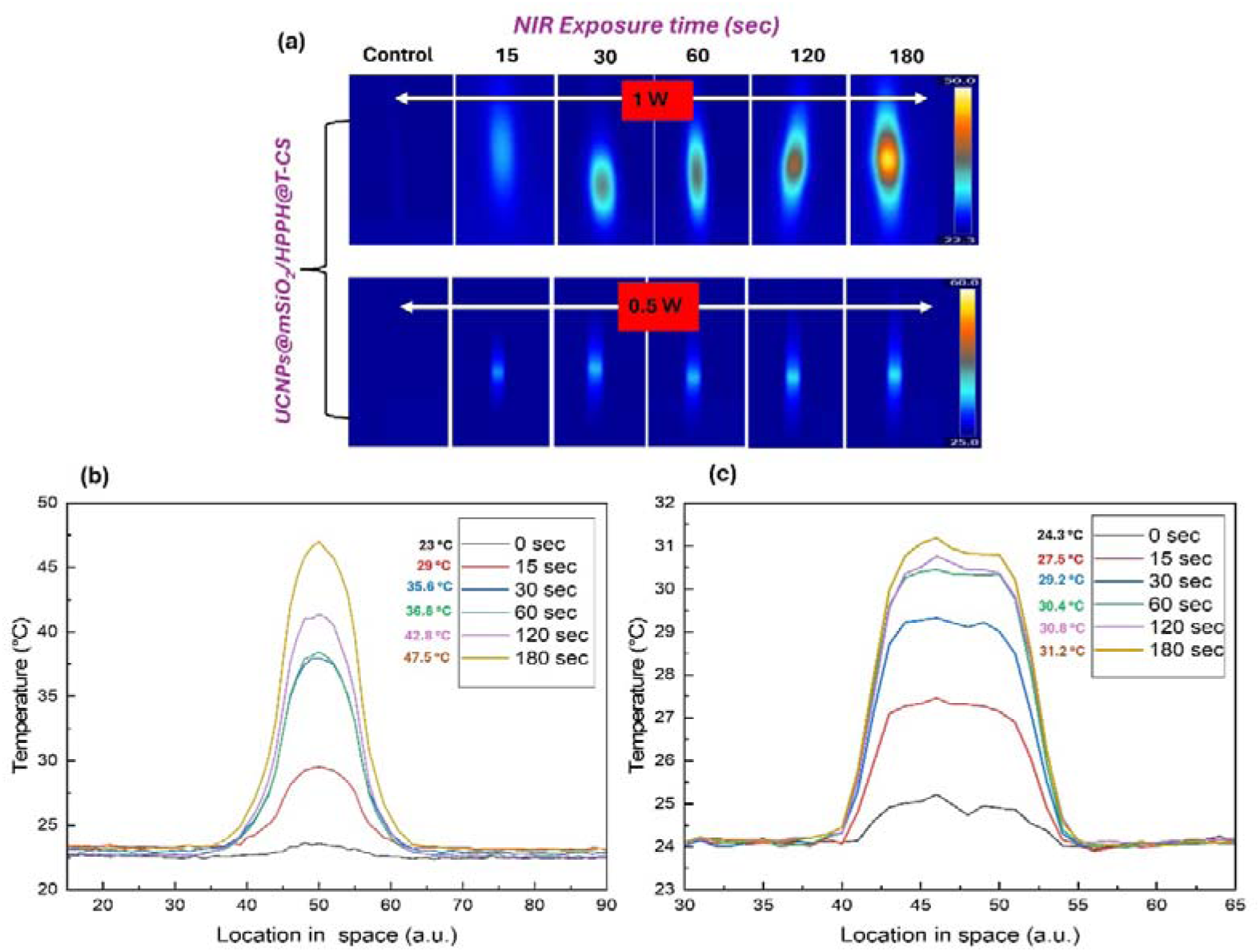
Evaluation of the photothermal heating behavior of the NF upon 980 nm laser irradiation. (a) Photothermal IR thermographs of the NF under 980 nm laser irradiation at 1 W; 1.27 W.cm^-2^ (top panel) and 0.5 W; 0.63 W.cm^-2^ (bottom panel) over various exposure times, demonstrating controlled, power-dependent temperature elevation. (b,c) Temperature elevation profiles of the NF under 980 nm laser irradiation over 180 s at irradiation powers of 1 W (b) and 0.5 W (c).

##### Optimized Power Screening (0.5 W (0.63 W.cm^-2^))

To suppress photothermal contributions, the laser power was reduced to 0.5 W, and temperature evolution was reassessed under identical conditions. At this lower power density, temperature increases were minimal and largely time-independent, with measured values of 27.5 °C (15 s), 29.2 °C (30 s), 30.4 °C (60 s), 30.8 °C (120 s), and a maximum of 31.2 °C at 180 s (Figure 2c). Importantly, temperatures remained well below physiological levels throughout the entire irradiation period, confirming that photothermal heating was negligible at this power. These acellular measurements define a safe operational window for PDT and demonstrate that UCNPs can be excited at 980 nm without inducing hyperthermia when appropriate laser parameters are selected.

#### 2.2.4. Evaluation of Acellular Singlet Oxygen and Hydroxyl Radical Generation Under PDT and SDT

The acellular ROS generation capability of UCNPs@mSiO₂/HPPH@TCS was systematically evaluated under PDT and SDT conditions. ¹O₂ Production was quantified using DPBF as a selective probe (Figure S12 (a, b)), while hydroxyl radical (^•^OH) generation associated with peroxidase-like (POD) catalytic activity was assessed using methylene blue degradation (Figure S13 (a, b)).

##### Singlet Oxygen Generation

A marked increase in ^1^O_2_ production was observed upon both photodynamic and sonodynamic activation of the NF, confirming the dual-modality ROS generation capability of the platform. The mean absorbance change (%) of ¹O₂ generation under PDT increased from 22.7 % at dose 1 to 31% at dose 2, reaching 36% at the highest irradiation dose (dose 3). This trend reflects efficient excitation of HPPH mediated by up-converted emissions from UCNPs under 980 nm laser irradiation. Similarly, SDT induced ¹O₂ generation with values of 6.2%, 9%, and 13.6% for doses 1, 2, and 3, respectively (Figure 3a). PDT consistently generated higher ^1^O₂ levels than SDT across all three doses, indicating that both modalities effectively activate the UCNPs@mSiO₂/HPPH@TCS nanoplatform, with PDT providing the advantage of greater singlet oxygen production due to the predominance of the type II photochemical mechanism, which efficiently converts molecular oxygen to ^1^O₂ under NIR laser irradiation. ^[35]^

**Figure 3.**
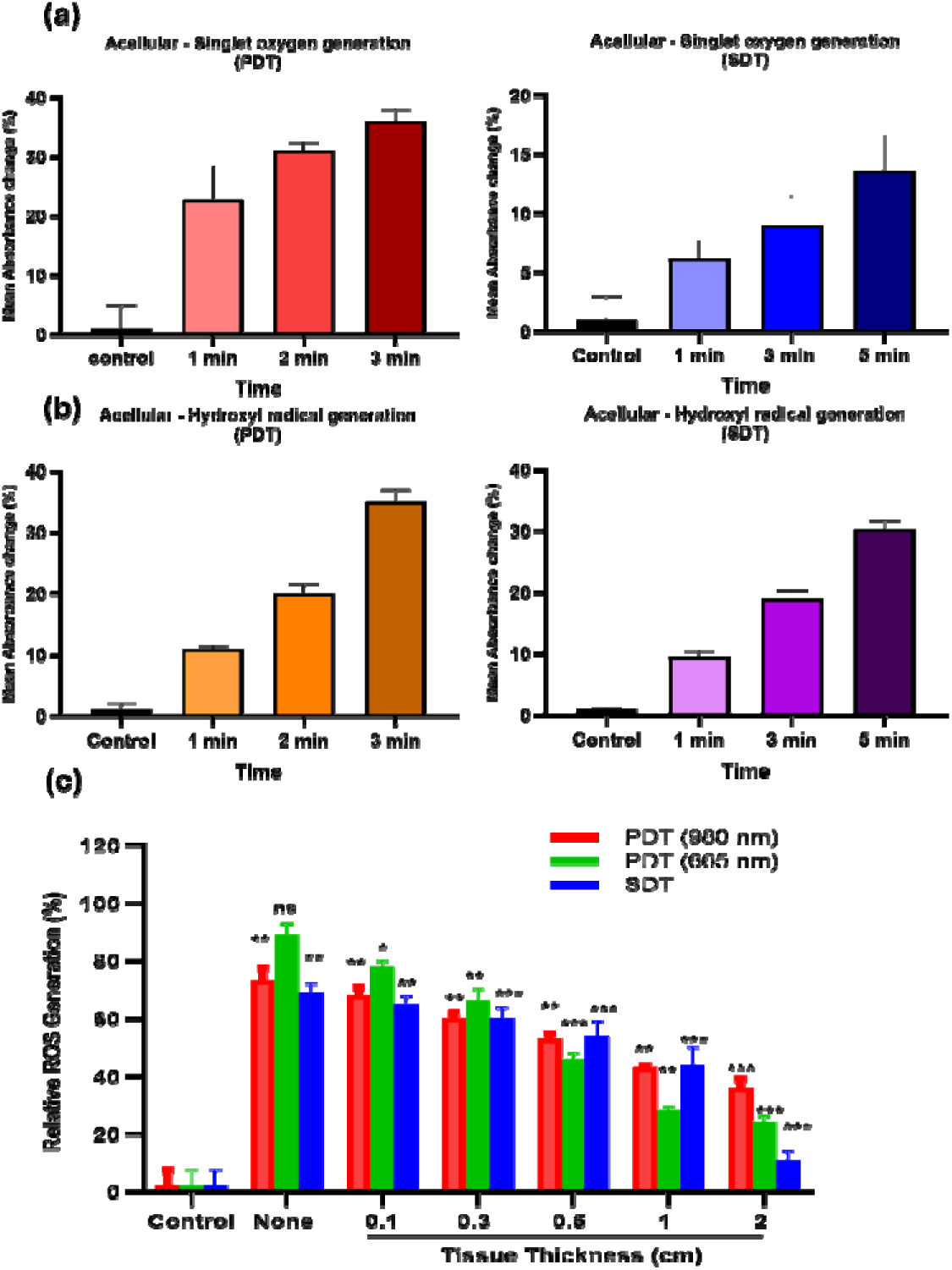
Acellular singlet oxygen and hydroxyl radical generation under PDT and SDT, along with evaluation of tissue depth penetration of the NF under 980 nm (0.63 W.cm^-2^) and 665 nm (0.63 W.cm^-2^) lasers and ultrasound waves. (a) Quantitative evaluation of singlet oxygen (^1^O₂) generation using DPBF degradation under PDT and SDT conditions. The efficiency of ROS production was expressed as the percentage of probe degradation calculated using the formula: [(A_0_ – A_t_) / A_0_] x 100. Under PDT irradiation, a progressive increase in DPBF degradation was observed with increasing exposure time, indicating controllable light-triggered ^1^O₂ generation. Similarly, SDT treatment induced time-dependent probe degradation during ultrasound exposure. However, PDT produced significantly higher singlet oxygen levels, reflected by the greater mean absorbance change (%) compared with SDT at corresponding time points. Negligible changes were observed in the control groups without activation, confirming the stability of the system in the absence of NIR irradiation or ultrasound stimulation. Data are presented as mean ± SD (n = 3). (b) Quantitative evaluation of hydroxyl radical (^●^OH) generation using methylene blue (MB) degradation under photodynamic (PDT) and sonodynamic (SDT) conditions. The efficiency of ^•^OH production was expressed as the percentage of probe degradation calculated using the formula: [(A_0_ – A_t_) / A_0_] x 100. Under PDT irradiation, a gradual increase in MB bleaching was observed with increasing exposure time, confirming the generation of hydroxyl radicals through the up-conversion-mediated photodynamic process. Similarly, SDT treatment resulted in a time-dependent increase in MB degradation during ultrasound exposure, demonstrating the sonocatalytic activity of the UCNPs@mSiO₂/HPPH@TCS nanoplatform. Control experiments showed minimal probe degradation, indicating that effective ^•^OH production requires activation of the nanoplatform. Data are presented as mean ± SD (n = 3). (c) Comparative depth-dependent ROS generation in acellular tissue models. Relative ROS production of the NF was quantified via the bleaching of the DPBF probe (ΔA = A_0_ - A_t_) after the respective triggers (980 nm NIR (0.63 W. cm^-2^), 665 nm visible light (0.63 W. cm^-2^), and ultrasound) were passed through porcine/bovine muscle tissue of varying thicknesses (0.1-2.0 cm). The Control represents the NF-loaded probe without external stimulation, while the None (0 cm) group serves as the 100% reference for unattenuated ROS output. SDT exhibits superior penetration at extreme depths (2 cm) compared to PDT modalities, which suffer from significant photon attenuation. Data are presented as mean ± SD (n=3), asterisks indicate statistical significance (*p < 0.05, **p < 0.01, ***p < 0.001) and ns denotes non-significance (p > 0.05) compared to the 0 cm reference.

##### Hydroxyl Radical Generation and POD-like Activity

The peroxidase-like catalytic activity of the nanoplatform was evaluated by monitoring hydroxyl radical generation via methylene blue degradation. Under PDT conditions, ^●^OH production increased markedly with dose, rising from 11% at dose 1 to 20% at dose 2 and reaching 35% at dose 3. An almost similar trend was observed under SDT, with corresponding values of 9.5%, 19%, and 30.3% (Figure 3b). Notably, hydroxyl radical generation exhibited comparable efficiencies under both PDT and SDT across all doses, particularly at higher energy inputs. This similarity indicates that ^•^OH production is primarily driven by energy-induced activation of the UCNPs@mSiO₂/HPPH@TCS nanoplatform rather than being strongly dependent on the specific excitation modality. Under both photonic and acoustic stimulation, enhanced energy deposition facilitates ROS formation through indirect radical-generating pathways, leading to comparable hydroxyl radical yields. The pronounced dose-dependent increase further confirms that higher energy input amplifies hydroxyl radical generation efficiency without modality-specific limitations.

#### 2.2.5. Depth-Dependent Evaluation of ROS Generation Under PDT and SDT

To systematically compare the penetration efficiency of 980 nm NIR light (0.63 W.cm^-2^), 665 nm visible light (0.63 W.cm^-2^), and US, a depth-dependent acellular ROS assay was conducted using DPBF as a ROS-sensitive probe. Fresh beef tissue slices of increasing thickness (0–20 mm) were placed between the irradiation source and the nanoparticle-containing Transwell® insert to simulate increasing tissue depth (Figure S14, S15).

##### ROS Generation Under Direct and Shallow Tissue Exposure

ROS generation was first evaluated under direct exposure without tissue obstruction, where 665 nm PDT produced the highest ROS yield (89%), followed by 980 nm PDT (73%) and SDT (69%). The superior performance of 665 nm irradiation under shallow conditions is attributed to direct excitation of HPPH at its absorption maximum, whereas ROS generation under 980 nm irradiation depends on the up-conversion process of UCNPs. SDT-induced ROS arises from acoustic cavitation and sonoluminescence, yielding comparable ROS levels under direct exposure. Upon introduction of tissue layers (1-3 mm), ROS generation decreased for all modalities, with more pronounced attenuation observed for 665 nm PDT due to increased scattering and absorption of visible light. Upon introducing a 3 mm tissue barrier, the ROS generation efficiency was reduced to 66.2% for 665 nm PDT, while both 980 nm PDT and SDT saw a steeper decline to 60% of their initial yields. At intermediate depths (5-10 mm), ROS generation under 665 nm irradiation declined sharply (45.5% and 28.3%), whereas both 980 nm PDT and SDT maintained substantially higher ROS levels (∼54% at 0.5 cm and ∼44% at 10 mm), highlighting sustained activation due to the penetration capability of NIR and US. At the maximum tested depth of 20 mm, ROS production further declined across all treatments; however, 980 nm PDT retained the highest ROS yield (36%), outperforming 665 nm PDT (24.3%) and SDT (11%). The pronounced reduction observed for SDT at greater depths is likely due to acoustic energy attenuation and reduced cavitation efficiency resulting from tissue scattering and impedance mismatch (Figure 3c). It should be noted that DPBF primarily detects ¹O₂ and exhibits limited sensitivity toward other ROS species, such as ^•^OH and superoxide anions. Consequently, SDT-induced ROS levels may be underestimated; nevertheless, the observed depth-dependent trends remain valid for comparative evaluation.^[36]^ Overall, these findings demonstrate that while visible-light PDT is most effective under shallow conditions, UCNP-mediated 980 nm PDT and SDT exhibit superior tissue penetration, with 980 nm PDT showing the greatest ROS retention at tested tissue-mimicking depths, underscoring its potential for deep-tissue PDT applications. The demonstration of therapeutic activation through a 20 mm tissue thickness represents a stringent and biologically meaningful benchmark for deep-tissue treatment, substantially exceeding the penetration limits of conventional visible-light PDT, which is typically restricted to a few millimeters. This depth is clinically relevant for many localized and orthotopic solid tumors, including prostate and other soft-tissue malignancies, where extracorporeal, endoluminal, or image-guided activation strategies are feasible. Importantly, while a 20 mm thickness does not fully recapitulate all remote metastatic settings or complex anatomical barriers, it provides a rigorous translational framework for evaluating penetration-enhanced therapeutic modalities such as sonodynamic and NIR-activated nanotherapeutics. Together, these findings underscore the potential of the present platform to overcome a central limitation of conventional PDT and support its further development toward clinically deployable deep-tissue cancer therapies.

### 2.3 Cellular Studies

#### 2.3.1. Cellular Uptake Kinetics in PSMA-Positive and PSMA-Negative Cells

To determine the optimal incubation period for maximizing intracellular accumulation of the nanoplatform before photodynamic and sonodynamic treatments, we evaluated its time-dependent cellular uptake in both LNCaP (PSMA^+^) and PC-3 (PSMA^-^) cell lines. Cells were exposed to the UCNPs@mSiO₂/HPPH@TCS for 0 (control), 1, 3, 6, 15, and 24 h, followed by fluorescence imaging to quantify intracellular nanoparticle levels. Uptake was assessed by measuring the red-region fluorescence intensity of HPPH using a Texas Red filter, providing a direct measure of internalized nanoplatforms (Figure 4a).

**Figure 4.**
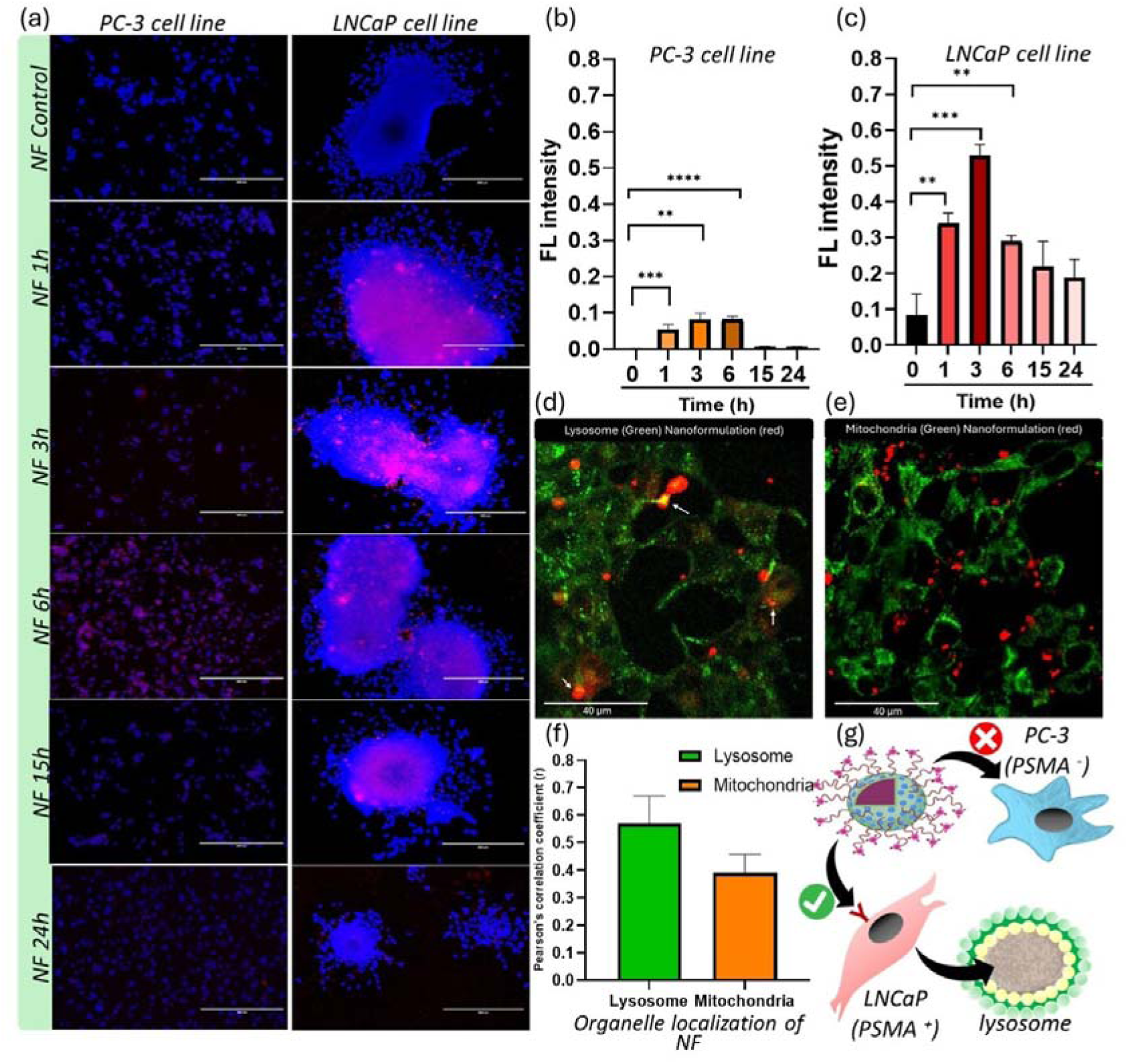
Receptor-Mediated Internalization and Subcellular Trafficking. (a) Representative time-dependent fluorescence microscopy images of PC-3 and LNCaP cells incubated with the NF over 24 hours. Nuclei are counterstained with DAPI (blue), and the red channel denotes the intrinsic fluorescence of the NF. (b, c) Quantitative analysis of intracellular uptake in PC-3 and LNCaP cells, demonstrating accelerated and significantly enhanced internalization in PSMA-positive LNCaP cells within 1–3 h post-treatment. (d,e) Representative confocal microscopy images illustrating the subcellular distribution of the NF relative to organelle-specific markers. The NF exhibits predominant colocalization with lysosomes, with negligible colocalization with mitochondria. (f) Quantitative evaluation of Pearson’s correlation coefficient (r) confirms high colocalization with lysosomal compartments compared to mitochondrial targets. (g) Schematic illustration of the receptor-mediated recognition and selective uptake mechanism of the PSMA-targeted UCNPs@mSiO_2_/HPPH@TCS NF in LNCaP cells versus PSMA^-^ PC-3 cells. Data are presented as mean ± SD (n=3). Statistical significance: ns, not significant; p ≤ 0.05 (*), p ≤ 0.01 (**), p ≤ 0.001 (***), and p ≤ 0.0001 (****).

Distinct uptake kinetics were observed between the two cell lines, with LNCaP cells showing rapid, PSMA-targeted endocytosis that maximized intracellular fluorescence by 3 h, whereas PC-3 cells displayed slower, non-specific minimum sustained levels at 1,3 and 6 h followed by a gradual reduction at later timepoints. The lower uptake in PC-3 cells aligns with their lack of PSMA expression and their non-specific nature, potentially due to enhanced permeability and retention (EPR) effects (Figure 4 (b, c)). These findings confirm that the targeting component of the nanoplatform substantially accelerates intracellular delivery in PSMA-expressing cells. Based on these kinetic profiles, incubation periods of 3 h for LNCaP cells and 6 h for PC-3 cells were selected as the optimal timepoints for subsequent PDT and SDT experiments, ensuring that therapeutic studies were conducted under conditions of maximal intracellular nanoparticle availability. Interestingly, fluorescence decreased at 6, 15, and 24 h, likely due to a combination of nanoparticle exocytosis, lysosomal degradation, and dilution among daughter cells during the fluorescent probe’s proliferation.^[37]^

#### 2.3.2. Subcellular Localization of UCNPs@mSiO₂/HPPH@TCS Nanoparticles

Next, we explored whether their trapping influences the subcellular distribution of nanoparticles in the lysosomes as well as electrostatic binding to mitochondria. In these experiments, LNCaP cells were incubated with UCNPs@mSiO₂/HPPH@TCS, stained with organelle-specific dyes, and imaged with confocal microscopy (Figure 4 (d, e)). After 3 hours of incubation, the nanoparticles exhibited efficient cellular uptake, as evidenced by strong intracellular fluorescence signals. Colocalization analysis revealed negligible overlaps between the NF signal and the mitochondrial marker, indicating that NF does not significantly bind to these organelles. In contrast, we observed a partial colocalization with LysoTracker-stained compartments, suggesting that a fraction of the internalized NF resided within lysosomes. This intracellular distribution pattern is consistent with the classical endocytic uptake pathway, in which NF enters the cell within endosomes and then merges with lysosomes before potential escape or redistribution to other organelles occurs. Single-cell colocalization analysis of UCNPs@mSiO₂/HPPH@TC NF with lysosomes and mitochondria was performed using Pearson’s correlation coefficient (Coloc2, Fiji). NF exhibited stronger lysosomal localization (mean R = 0.57 ± 0.16, n = 10 cells) compared to mitochondrial localization (mean R = 0.39 ± 0.21, n = 10 cells). Individual cell values ranged from 0.29 to 0.73 for lysosomes and 0.03 to 0.75 for mitochondria, indicating variable intracellular distribution (Figure 4 (f,g)).^[38]^ Importantly, lysosomal accumulation may be therapeutically relevant, as lysosomes provide an acidic microenvironment that can facilitate PS activation and ROS generation upon photodynamic or sonodynamic stimulation, potentially triggering lysosomal membrane permeabilization-mediated cell death pathways. Taken together, these results indicate that UCNPs@mSiO₂/HPPH@TCS NF are rapidly internalized by cancer cells and primarily processed through the endolysosomal pathway during the early uptake phase.

#### 2.3.3. Biocompatibility and Cytotoxicity Evaluation

The cytotoxicity of free HPPH and HPPH-loaded NF (HPPH@NF) was assessed in LNCaP (PSMA^+^) and PC-3 (PSMA^-^) PCa cell lines using a Cell Counting Kit-8 (CCK-8) cell viability assay. Cells were incubated with increasing concentrations of HPPH or HPPH@NF (1, 2, 3, 5, 10, and 20 µg. mL^-1^) for 24 h under dark conditions to evaluate the intrinsic biocompatibility of the formulations before SDT/PDT treatments. In LNCaP cells, both free HPPH and HPPH@NF demonstrated high biocompatibility across all tested concentrations, maintaining cell viability > 80%, indicating minimal dark toxicity and good physiological tolerance of the PSMA-targeted nanoplatform. In contrast, in PC-3 cells, viability remained above 80% at concentrations only up to 3 µg. mL^-1^ for both free HPPH and HPPH@NF, and at concentrations above 5 µg. mL^-1^ we observed a progressive decline in PC-3 viability, reaching < 40% at 20 µg. mL^-1^. The preservation of PC-3 cell viability at lower HPPH concentrations likely reflects the intrinsic resistance and elevated antioxidant capacity of androgen-independent CRPC cells. However, once intracellular HPPH levels exceed a critical threshold (≥ 5 µg. mL^-1^), oxidative stress surpasses cellular defense mechanisms, resulting in a progressive, dose-dependent loss of viability, culminating in marked cytotoxicity at higher concentrations. Based on these observations, a concentration of 3 µg. mL^-1^ was selected as the safe and physiologically compatible dose for all subsequent mechanistic and therapeutic assays in both LNCaP and PC-3 cell lines. This ensured that downstream SDT/PDT effects could be attributed to nanoparticle-mediated therapeutic mechanisms rather than dark cytotoxicity (Figure 5a).

**Figure 5.**
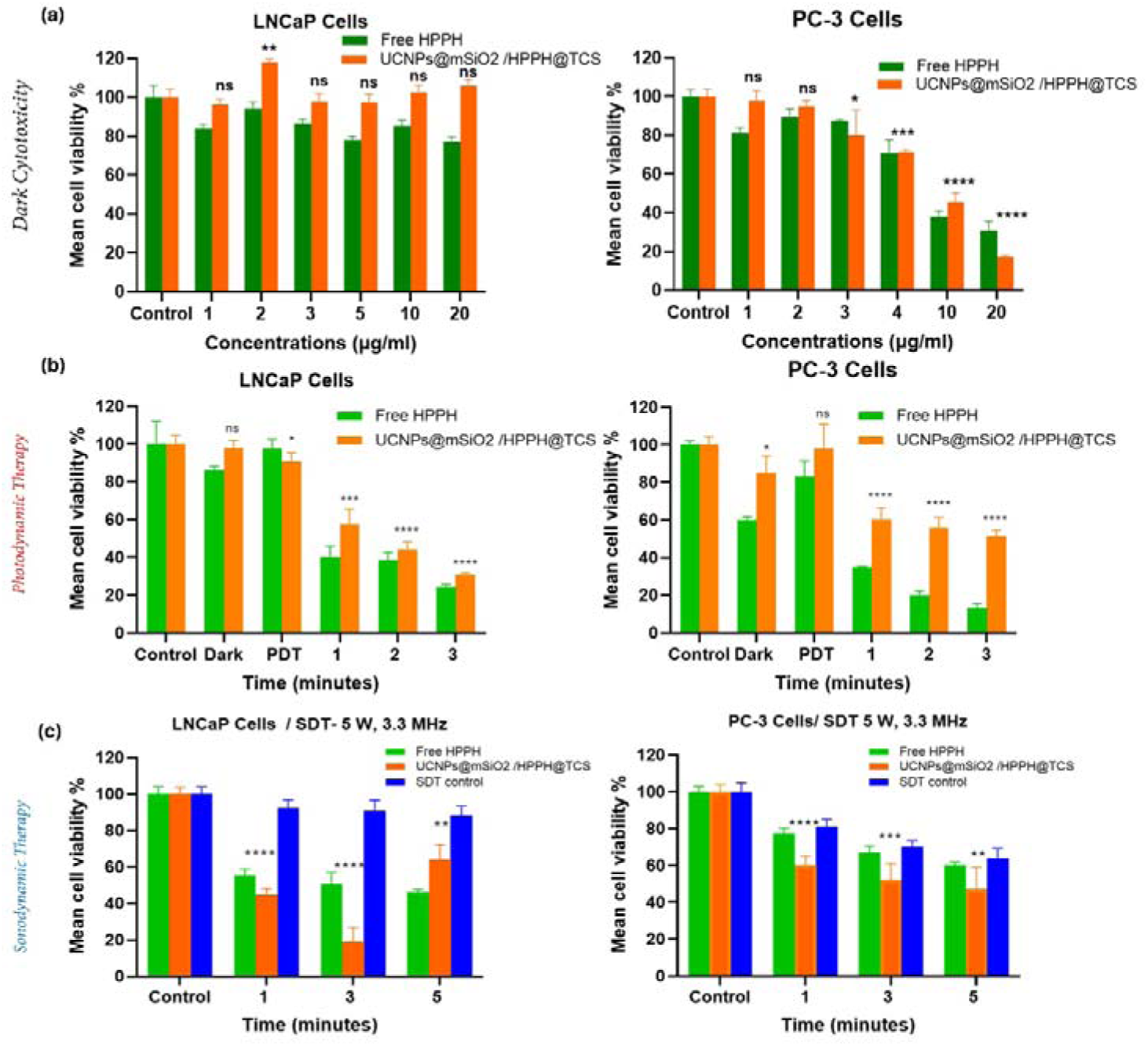
Targeted In Vitro Therapeutic Efficacy of NF in LNCaP and PC-3 cells. (a) Dark cytotoxicity evaluation of the NF in PC-3 and LNCaP cells (1–20 µg.ml-1), confirming high biocompatibility at the working dosage. (b) Comparative cell viability of PC-3 and LNCaP cells following PDT (980 nm (0.63 W.cm^-2^) irradiation for NF and 665 nm (0.63 W.cm^-2^) for free HPPH) at increasing fluences (37.8, 75.6, and 113.4 J.cm^-2^). (c) Therapeutic efficacy of SDT (3 MHz, 5 W) in PC-3 and LNCaP cells across various exposure times, comparing free HPPH, the targeted NF, and ultrasound-only controls. Data represent mean ± SD (n=3). Statistical significance: ns, not significant; p ≤ 0.05 (*), p ≤ 0.01 (**), p ≤ 0.001 (***), and p ≤ 0.0001 (****).

#### 2.3.4. PDT Cell Viability results in PC-3 and LNCaP Cells

Distinct therapeutic responses emerged as we combined NF with NIR irradiation. Our results in Figure 5b show, in PC-3 cells, PDT induced only a modest decrease in viability, with survival remaining near ∼50% even at the highest dose, consistent with limited NF internalization in PSMA^−^ cells and consequently reduced intracellular ROS generation. In contrast, PSMA^+^ LNCaP cells exhibited a pronounced and dose-dependent phototoxic response, with cell viability decreasing to below 20 % at 113 J·cm⁻² (Figure 5b). This enhanced therapeutic efficacy in LNCaP cells reflects PSMA-mediated NF uptake facilitated by the targeted chitosan coating, resulting in higher intracellular HPPH accumulation and amplified ROS production upon UCNP-mediated excitation. Together, these findings demonstrate that the observed cytotoxicity arises from PSMA-specific photodynamic activation rather than nonspecific NF or irradiation effects, underscoring the critical role of molecular targeting in achieving selective therapeutic outcomes.

#### 2.3.5. SDT Cell Viability results in PC-3 and LNCaP Cells

A notable feature of the SDT response was the emergence of a biphasic viability profile in PSMA^+^ LNCaP cells at 3 MHz frequency and high acoustic intensity (5 W.cm^-2^). Short sonication durations (1-3 min) produced the expected enhancement in SDT-induced cytotoxicity, consistent with efficient nanoparticle activation and intracellular ROS generation. However, extending exposure to 5 min paradoxically resulted in partial recovery of cell viability (Figure 5c). This non-monotonic response contrasts with the behavior observed in PSMA^-^ PC-3 cells, where cell viability decreased progressively with increasing sonication time and intensity, reflecting a primarily energy-dependent SDT response that is independent of receptor-mediated nanoparticle uptake. The divergent behavior between the two cell lines can be rationalized by the PSMA-targeted nature of the UCNPs@mSiO₂/HPPH@TCS NF. In LNCaP cells, therapeutic efficacy is strongly coupled with increased uptake due to PSMA-mediated internalization of the NF. At elevated acoustic power and prolonged exposure, US-induced mechanical effects, including microstreaming, shear stress, and transient cavitation, can disrupt ligand-receptor interactions, alter the conformation or integrity of the chitosan targeting layer, or promote partial desorption of the NF coating, which has been reported by others as well.^[39]^ These effects would reduce effective intracellular NF accumulation during extended sonication, thereby limiting ROS production and attenuating SDT efficacy at the longest exposure time. In contrast, PSMA^-^ PC-3 cells, which rely on non-specific NF uptake, are less sensitive to such targeting-related perturbations and therefore exhibit a monotonic, dose-dependent SDT response driven primarily by acoustic energy deposition.

#### 2.3.6. Comparative Analysis of PDT and SDT-Induced ROS Production in PSMA^-^ & PSMA^+^ Prostate Cancer Cells

The intracellular ROS generation mediated by UCNPs@mSiO₂/HPPH@TCS was evaluated in PSMA^-^ (PC-3) and PSMA^+^ (LNCaP) PCa cells under PDT and SDT conditions. The ROS production was quantified at three different time points following treatment, using the pre-determined optimal exposure parameters: 0.5 W (0.63 W.cm^-2^) for 980 nm laser irradiation (PDT) and 5 W.cm^-2^, 3 MHz US (SDT). Results shown in Figure 6 (a,b) demonstrate markedly enhanced ROS generation in PSMA-expressing LNCaP cells compared with PSMA^-^ PC-3 cells. Under SDT conditions, ROS intensity increased progressively with dose, reaching a maximum value of 2.62 at the highest exposure level. PDT activation also produced a clear dose-dependent ROS response, with fluorescence intensities increasing from 0.13 to 1.20 across the tested dose range. Although SDT produced higher absolute ROS signals compared to PDT, both activation modalities demonstrated efficient intracellular ROS generation in PSMA^+^ LNCaP cells. These results confirm that the UCNPs@mSiO₂/HPPH@TCS nanoplatform effectively mediates ROS production under both optical and acoustic stimulation, supporting its potential for multimodal therapeutic activation. Furthermore, SDT induced greater ROS production than PDT across all time points in both cell lines. This observation likely reflects the broader spectrum of ROS generated under SDT, including ^1^O₂, ^•^OH, and superoxide anions (O₂^•−^), whereas PDT predominantly generates singlet oxygen via type II photochemical pathways. ^[40]^ The time-course analysis demonstrated a progressive increase in ROS generation under both PDT and SDT, with SDT consistently producing higher ROS levels at each measured interval (Figure S12, S13). These findings indicate that the UCNPs@mSiO₂/HPPH@TCS platform can induce robust oxidative stress in PSMA^+^ PCa cells, with SDT providing a more pronounced ROS response than PDT under the optimized treatment conditions.

**Figure 6.**
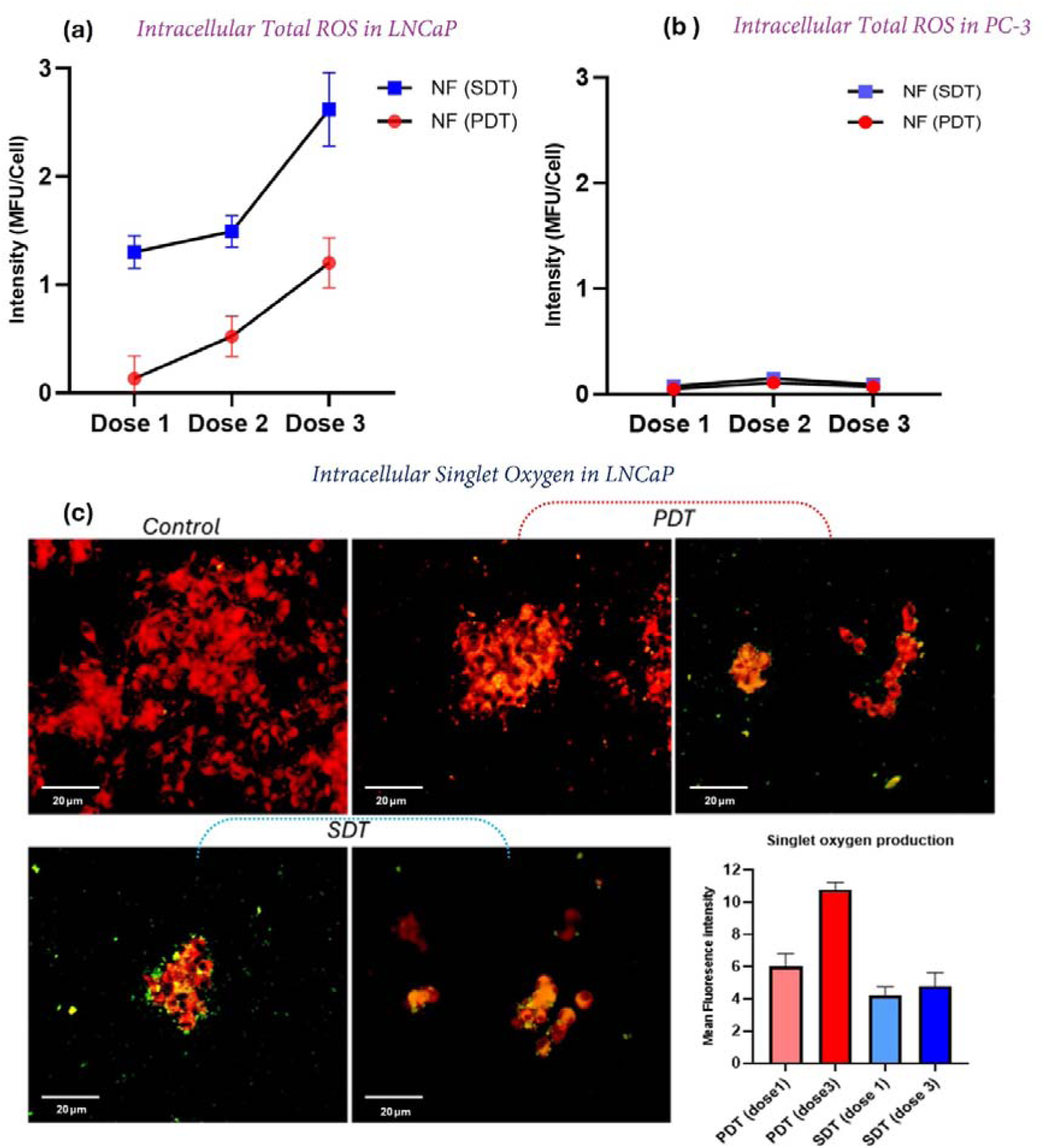
Quantitative analysis of total intracellular ROS and Singlet Oxygen generation induced by the NF under PDT and SDT. (a,b) ROS levels were assessed via mean fluorescence intensity (MFI) as a function of light dose (37.8-113.4 J.cm-2) and sonication time (1-5 min) in LNCaP and PC-3 cells respectively. (c) Representative fluorescence microscopy images of intracellular singlet oxygen (^1^O2) production in LNCaP cells using SOSG (green fluorescence) following NF-mediated PDT and SDT. The images demonstrate dose-dependent and modality-dependent ^1^O2 generation. Data represent mean ± SD (n=3). Statistical significance: ns, not significant; p ≤ 0.05 (*), p ≤ 0.01 (**), p ≤ 0.001 (***), and p ≤ 0.0001 (****).

#### 2.3.7. Cellular ¹O₂ Dynamics in PSMA^+^ LNCaP Cells Visualized by SOSG

Intracellular ¹O₂ generation in PSMA^+^ LNCaP cells was visualized using SOSG following NF-mediated PDT and SDT at two dose levels. Overall, PDT consistently produced higher levels of ¹O₂ than SDT at both doses, with a pronounced dose-dependent increase observed under photodynamic activation. Specifically, PDT induced a substantial rise in SOSG fluorescence intensity from 6 at the lower dose (Dose 1) to 10.78 at the higher dose (Dose 3), indicating efficient ¹O₂ generation with increasing light energy input (Figure 6c). This behavior is consistent with the classical Type II photochemical mechanism of PDT, in which enhanced photon absorption by the PS promotes intersystem crossing to the triplet state and subsequent energy transfer to molecular oxygen, resulting in elevated ¹O₂ production.^[41]^ In contrast, SDT resulted in lower overall ¹O₂ levels and exhibited minimal dose dependence, with mean fluorescence intensities of 4.2 and 4.8 for Dose 1 and Dose 3, respectively. The limited increase in SOSG signal at higher US dose suggests that, under the applied conditions, SDT-induced cytotoxicity may not be primarily driven by ¹O₂ generation. Instead, US activation is known to generate a broader spectrum of reactive oxygen species through acoustic cavitation, sonoluminescence, and radical-mediated pathways, including ^•^OH and superoxide anions, which are not efficiently detected by SOSG. Consequently, the relatively flat ¹O₂ response across SDT doses likely reflects both mechanistic differences in ROS generation and probe selectivity limitations, rather than a lack of biological activity. Collectively, these findings highlight a fundamental mechanistic distinction between PDT and SDT in PSMA^+^ PCa cells. While PDT demonstrates strong, dose-dependent singlet oxygen production, SDT appears to elicit oxidative stress through non-¹O₂-dominant pathways, which may contribute to its therapeutic efficacy through complementary mechanisms.^[40]^ This divergence underscores the value of integrating both modalities within a single nanoplatform, enabling differential ROS engagement and potentially enhancing overall anticancer efficacy in PCa.

#### 2.3.8. Effects on Tumor Progression, Inflammatory, and Apoptotic Gene Expression

Given the central role of proliferative signaling, angiogenesis, apoptosis, and inflammation in PCa progression and therapeutic resistance, we evaluated treatment-induced modulation of key molecular regulators within these pathways following NF, PDT, and SDT treatments (Figure 7). The gene panel encompassed tumor progression and survival regulators (AKT1, MYC), prostate cancer-associated markers (KLK3), angiogenic mediators (VEGFA), apoptotic regulators (BCL2, BCLX), executioner caspases (CASP3, CASP7), and the pro-inflammatory cytokine IL-1β. Overall, both PDT and SDT elicited distinct, treatment-dependent transcriptional responses relative to NF treatment alone, reflecting fundamentally different stress modalities and downstream cellular adaptations.

**Figure 7.**
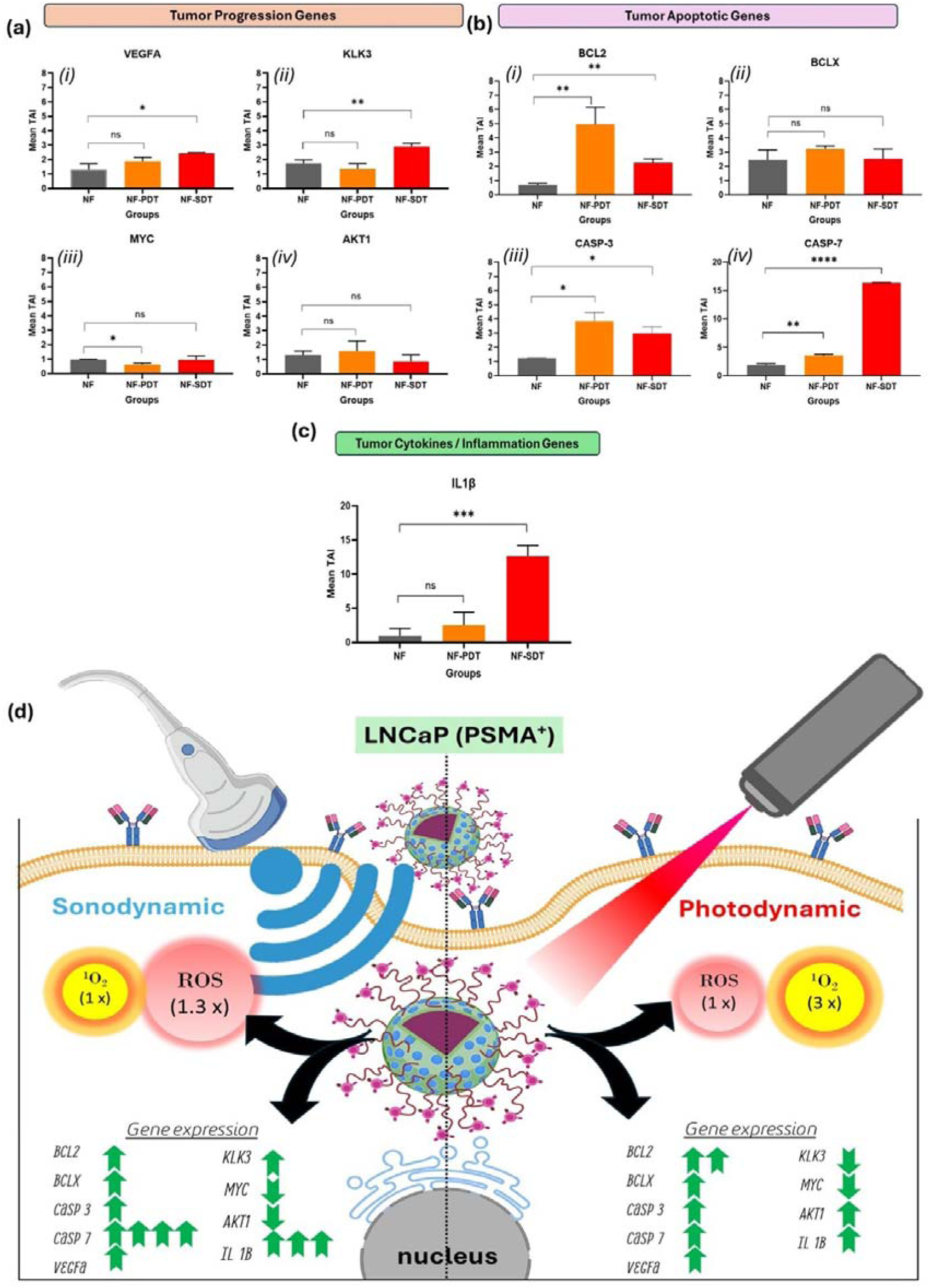
Comparative gene expression profiling of angiogenesis, proliferation, apoptosis, and inflammatory responses induced by PDT and SDT. **(a)** Regulators of Angiogenesis and Tumor Progression: Expression profiles of VEGFA, KLK3, MYC, and AKT1. Note the distinct angiogenic rebound (VEGFA) across both therapies and the marked divergence in KLK3 and MYC signaling, where PDT favors acute proliferative rescue (high MYC) while SDT drives biomechanical stress responses (KLK3). **(b)** Apoptotic and Inflammatory Signaling: Expression of anti-apoptotic factors (BCL2, BCLX), executioner caspases (CASP3, CASP7), and **(c)** the pro-inflammatory cytokine IL-1β. While PDT triggers robust early anti-apoptotic defenses (BCL2), SDT effectively bypasses these survival signals to drive a dramatic induction of CASP7 and IL-1β, signaling a shift toward irreversible cell death and immunogenic stress. **(d)** Schematic illustration of the mechanism of PSMA⁺ prostate cancer cell death triggered by the NF under photo and sonodynamic therapy.

Our initial determination was on tumor progression genes, of which the VEGFA gene expression was increased following both PDT and SDT compared with NF treatment alone (Figure 7a (i)). PDT resulted in a ∼ 1.5-fold elevation in VEGFA expression; however, this change did not achieve statistical significance (TAI: 1.90 vs. 1.30; p = ns). In contrast, SDT elicited a significant induction of VEGFA, with expression level increasing by ∼ 1.7–1.8-fold over NF controls (TAI ∼2.25; p < 0.05). The modest non-significant increase in VEGFA following PDT is consistent with a secondary consequence to ¹O₂-mediated oxidative stress, a defining feature of PDT that promotes localized hypoxia.^[42]^ Hypoxia-driven stabilization of hypoxia-inducible factors and subsequent activation of pro-angiogenic signaling pathways are well-documented adaptive responses to photo-oxidative injury.^[43]^ By comparison, the more pronounced VEGFA induction following SDT aligns with an injury-associated tissue stress and wound-healing response.^[44]^ SDT involves US-mediated acoustic cavitation and ROS generation, processes that can induce transient hypoxia, cellular disruption, and stromal activation within the tumor microenvironment.^[43]^ Together, these data suggest PDT and SDT overlapping but mechanistically distinct angiogenic responses, reflecting early compensatory signaling rather than sustained angiogenic reprogramming. Nonetheless, even transient VEGFA upregulation may represent a potential adaptive escape pathway, supporting the rationale for incorporating anti-angiogenic or tumor microenvironment–modulating strategies to mitigate therapy-induced resistance in future translational applications. In this panel, we next examined the expression of KLK3, a PCa-associated, androgen-responsive gene (Figure 7a (ii)). KLK3 expression was modestly reduced following PDT compared with NF treatment alone (TAI: 1.35 vs. 1.76); however, this difference did not reach statistical significance. In contrast, SDT induced a statistically significant increase in KLK3 expression (TAI ∼2.91; p < 0.01), corresponding to a ∼1.7-fold elevation relative to NF controls. The pronounced induction of KLK3 following SDT suggests engagement of stress-responsive transcriptional programs that extend beyond those activated by PDT. US-mediated membrane perturbation, cytoskeletal deformation, and mechanotransduction signaling have been reported to modulate androgen receptor-associated transcriptional activity, which may contribute to the observed increase in KLK3 expression.^[45]^ Collectively, these findings are consistent with SDT eliciting a biomechanical stress signature distinct from predominantly photo-oxidative injury.

Following this, we analyzed the expression of MYC and AKT1, two key regulators of proliferative and survival signaling (Figure 7a (iii), 7a (iv)). MYC expression, a master regulator of cellular proliferation and metabolic adaptation, did not differ significantly following either PDT or SDT (PDT: TAI = 1.24, p = ns; SDT: TAI = 1.13, p = ns). Similarly, AKT1 transcript levels showed no significant change following PDT or SDT when compared with NF treatment alone, mirroring the absence of transcriptional modulation observed for MYC. Although MYC downregulation has been reported in select experimental systems following photodynamic intervention, our findings are consistent with studies demonstrating that PDT-induced cytotoxicity is predominantly mediated by ROS-driven apoptotic or necrotic injury, rather than transcriptional repression of oncogenic regulators.^[46]^ Accordingly, the therapeutic effects observed in this system are more likely attributable to acute oxidative cellular damage than to modulation of MYC-dependent transcriptional programs.^[47]^ The conserved transcriptional stability of MYC and AKT1 further suggests that PDT and SDT do not primarily exert their antitumor effects through sustained gene-level reprogramming of oncogenic signaling pathways. Instead, consistent with the established mechanisms of both modalities, therapeutic efficacy is likely mediated by acute, post-translational disruption of signaling networks, including oxidative stress and mechanotransduction-associated modulation of kinase activity. Collectively, these findings support a model in which PDT and SDT induce acute cytotoxic stress, with therapeutic efficacy driven largely by rapid, non-transcriptional disruption of survival signaling, accompanied by short-lived compensatory transcriptional responses in select microenvironment-responsive genes, such as VEGFA.

Analysis of apoptosis-related genes revealed distinct and temporally layered regulation of survival and death pathways (Figure 7b). Anti-apoptotic BCL2 expression was significantly elevated following PDT (∼4.95-fold, p<0.01), whereas SDT induced a more modest increase (∼2.25-fold, p=ns). This finding indicates that PDT triggers a strong acute anti-apoptotic stress response, likely reflecting early cellular attempts to counteract oxidative damage. The comparatively lower induction of BCL2 under SDT suggests reduced engagement of survival signaling, consistent with more severe or less reversible cellular injury (Figure 7b (i)). In contrast to the increased BCL-2 gene expression observed following PDT and SDT, BCL-X expression remained unchanged relative to NF treatment alone, indicating selective transcriptional engagement within the anti-apoptotic BCL-2 family. This differential response suggests that PDT and SDT-induced stress preferentially activate compensatory BCL-2-mediated survival signaling at the transcriptional level, while baseline BCL-X expression is maintained to preserve mitochondrial integrity without requiring further upregulation (Figure 7b (ii)). Together, the combination of elevated BCL-2 expression and stable BCL-X levels implies that anti-apoptotic buffering capacity is retained, potentially limiting a sufficient shift toward pro-apoptotic signaling and contributing to partial resistance to PDT and SDT-induced cytotoxic stress. Importantly, this early pro-survival signaling does not contradict apoptotic activation but rather reflects the well-established phenomenon in which cancer cells transiently upregulate anti-apoptotic defenses before succumbing to irreversible damage.^[48]^ Consistent with this interpretation, executioner caspases were robustly induced, with the most pronounced effects observed following SDT. CASP3 expression increased significantly from a TAI of 1.24 with NF treatment alone to 3.85 following PDT (∼ 3.1-fold; p < 0.01) and to 2.98 following SDT (approximately 2.4-fold; p < 0.01). Notably, CASP7 exhibited a marked and highly significant induction following SDT (TAI = 16.44; p < 0.001), corresponding to an approximately 8.7-fold increase relative to NF controls. The magnitude of CASP7 upregulation following SDT is consistent with strong engagement of execution-phase apoptotic programs. Prior studies have shown that US-mediated cavitation and associated cellular stress can promote mitochondrial and lysosomal destabilization, processes that facilitate caspase activation and irreversible commitment to cell death. ^[49, 50]^ Collectively, these findings support efficient activation of apoptotic machinery following SDT, potentially exceeding that observed with PDT (Figure 7b (iii), 7b (iv)).

Finally, expression of the pro-inflammatory cytokine IL-1β was minimally increased following PDT (TAI = 2.52; ∼2.7-fold), and dramatically elevated following SDT (TAI = 12.65; ∼13.5-fold vs NF). This pronounced SDT-induced inflammatory response consists of immunogenic cell stress, damage-associated molecular pattern (DAMP) release, and extensive cellular disruption (Figure 7c). Such inflammatory signaling may be advantageous for promoting anti-tumor immune activation and supports the potential integration of SDT with immunomodulatory or immune-checkpoint-based strategies.^[51]^

Taken together, these results demonstrate that while both PDT and SDT activate compensatory pro-survival pathways, SDT more effectively drives tumors toward irreversible apoptosis and immunogenic inflammation, whereas PDT elicits stronger early survival signaling that may transiently counteract cell death. These mechanistic distinctions provide a strong rationale for therapy-specific combination strategies and highlight SDT as a particularly potent modality for overcoming adaptive resistance mechanisms in prostate cancer (Figure 7d).

#### 2.3.9. Size-Dependent Therapeutic Response of NF-Mediated SDT and PDT in LNCaP Tumor Spheroids

To interrogate size-dependent therapeutic responsiveness, LNCaP spheroids matured for 6 and 14 days were subjected to SDT or PDT activation following incubation with the NF (Figure 8a). These two maturation stages yielded compact spheroids of distinct diameters (6 day: ∼1310-1407 μm; 14 day: ∼2289-2668 μm), enabling evaluation of treatment penetration and efficacy across increasing tumor-like mass and architectural complexity. Baseline imaging was first performed on NF-labeled spheroids in the absence of the pre-apoptotic marker pSIVA to confirm structural integrity and fluorescence distribution. Subsequent experiments incorporated pSIVA to dynamically monitor early apoptotic membrane inversion following SDT or PDT activation. Time-resolved imaging (0, 1, 3, 6, 12, 24, 36, and 48 h) enabled quantitative and morphological assessment of treatment-induced spheroid degradation (Figure 8 b, c).

**Figure 8.**
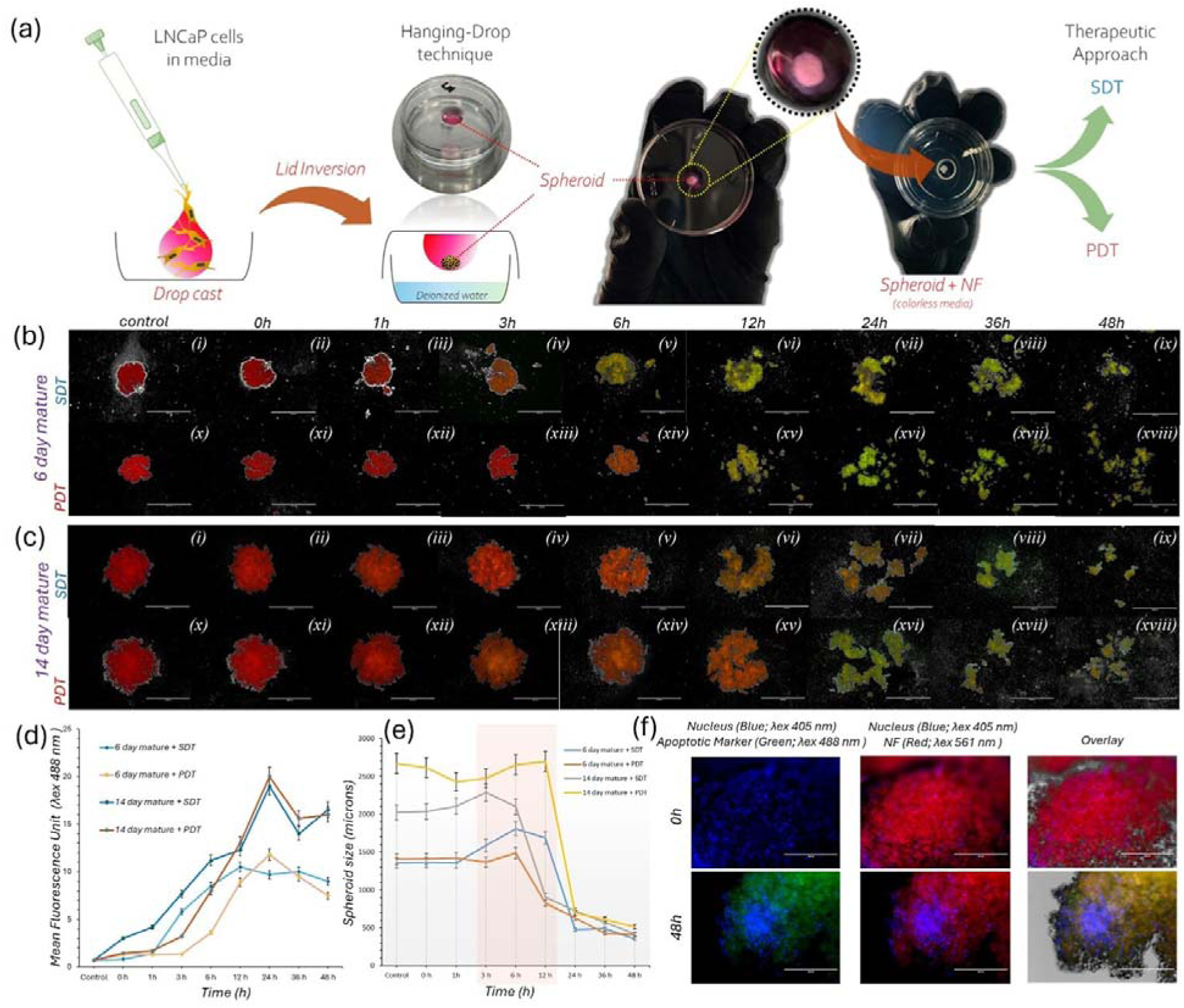
Spheroid maturation and therapeutic evaluation of UCNPs@mSiO_2_ /HPPH@TCS. (a) Schematic of LNCaP spheroid generation *via* hanging-drop technique and subsequent therapeutic approach (SDT vs. PDT). (b–c) Time-course fluorescence imaging of (b) 6-day and (c) 14-day mature spheroids following treatment, showing structural disintegration over 48 h. (d) Mean fluorescence intensity (λ_ex_ = 488 nm) over time, quantifying treatment-induced signaling. (e) Changes in spheroid diameter (microns) post-treatment; the shaded area indicates the onset of rapid disintegration. (f) Fluorescence images at 0 and 48 h showing nuclei (blue), nanoformulation (red), and apoptotic markers (green), with merged overlays.

Under SDT conditions, apoptosis was initiated as early as 3 h in both 6-day (∼1310 μm) and 14-day (∼2289 μm) spheroids, accompanied by early architectural destabilization and fragmentation of the tumoroid structure. In contrast, PDT-treated spheroids (6 day: ∼1407 μm; 14 day: ∼2668 μm) exhibited a delayed apoptotic response, with clear pSIVA activation emerging at ∼6 h and structural fragmentation occurring predominantly between 6-12 h. These findings indicate that NF-mediated cytotoxicity is triggered more rapidly under sonodynamic activation compared to photodynamic exposure, independent of spheroid size (Figure 8d).

Morphologically, fragmentation initiated from both peripheral and core regions of the spheroids, suggesting effective NF penetration and relatively uniform intratumoral distribution within the 3D mass. For both SDT and PDT, pronounced structural disintegration was evident between 12 and 24 h, consistent with progressive loss of cell-cell adhesion and collapse of multicellular integrity, potentially associated with downregulation of adhesion-associated proteins such as ICAM-1 in PSMA⁺ LNCaP cells.

Quantitative spheroid size analysis (Figure 7e) corroborated these observations. Under SDT, 6-day spheroids decreased from 1355 μm to 354 μm, while 14-day spheroids contracted from 2033 μm to 421 μm by 48 h. Similarly, PDT treatment reduced 6-day spheroids from 1407 μm to 413 μm and 14-day spheroids from 2668 μm to 514 μm. Notably, a transient increase in apparent spheroid diameter between 3-12 h was observed across treatment groups, attributable to internal core fragmentation and outward expansion before complete structural collapse.

A notable reduction in red fluorescence intensity was observed in the PDT-treated groups at later time points (Figure 8b (xvi-xviii) and 8c (xvi-xviii)), a result consistent with the progressive photobleaching of the HPPH photosensitizer within the NF system under continuous light activation. In contrast, high-resolution (20×) imaging of the SDT-treated spheroids at 0 and 48 h demonstrated extensive and irreversible apoptosis (Figure 8f). This therapeutic response was characterized by pronounced chromatin condensation (pyknosis) and nuclear accumulation within the post-treatment spheroid remnants. Significantly, the ultrasound-mediated activation in the SDT group appeared to bypass the photobleaching limitations observed in the PDT-mediated cohorts, resulting in more sustained and uniform structural degradation over the 48-h period. Collectively, these results demonstrate that SDT induces a more rapid and penetrative apoptotic response compared to PDT across both intermediate and large spheroid models, supporting the enhanced therapeutic kinetics of sonodynamic activation in densely packed 3D tumor architectures.

## 3. Conclusion

This work reports a UCNPs@mSiO₂/HPPH@TCS NF that enables a direct mechanistic comparison of penetration-enhanced PDT and SDT for PCa. By integrating up-conversion-mediated NIR activation, mesoporous silica-based PS loading, and PSMA-directed chitosan targeting within a single nanoarchitecture, we address the intrinsic depth limitations of conventional PDT. Response-surface-guided optimization enabled precise control of ROS generation while minimizing photothermal contributions, demonstrating that although visible-light PDT is effective at shallow depths, UCNPs-mediated NIR-PDT and SDT provide superior tissue penetration, with SDT generating higher total ROS through multimodal radical pathways. The physical advantages of our NF translated into enhanced biological efficacy in PSMA⁺ cells, including targeted uptake, lysosomal localization, elevated intracellular ROS generation, apoptosis induction, and activation of pro-apoptotic gene expression, resulting in rapid therapeutic responses in 3D tumor spheroids. Compared with PDT, SDT induced earlier apoptosis, more uniform spheroid penetration, and faster structural degradation, consistent with superior NF-mediated ROS delivery and intratumoral distribution.

Collectively, our findings establish the NF as a highly promising platform for optically inaccessible prostate tumors, demonstrating size-independent penetration and rapid apoptotic induction. Building upon these results, our future research will focus on the synergistic integration of these modalities into a unified photo-sonodynamic therapy (PSDT) regimen. Furthermore, we aim to evaluate the therapeutic efficacy and systemic biocompatibility of this targeted NF in vivo using PSMA^+^ tumor xenograft models. This approach seeks to transition our mechanistic benchmarking into a clinically relevant strategy for the precision treatment of dense, deep-seated 3D tumor architectures.

## 4. Experimental Section/ Methods

### 4.1. Materials

ErCl₃·6H₂O (99.99%), YbCl₃·6H₂O (99.99%), YCl₃·6H₂O (99.99%), 3-aminopropyltriethoxysilane (APTES), and tetraethyl orthosilicate (TEOS, 99.9%) were purchased from Sigma-Aldrich. Sodium hydroxide (NaOH, 96%), ammonium fluoride (NH₄F, 98%), ammonium hydroxide (NH₄OH), ammonium nitrate (NH₄NO₃), methanol (CH₃OH, 99.5%), 1-ethyl-3-(3-dimethylaminopropyl) carbodiimide (EDC), and N-hydroxysuccinimide (NHS) were also obtained from Sigma-Aldrich. Oleic acid (OA, 90%) and 1-octadecene (ODE, 90%) were purchased from Sigma-Aldrich. Phosphate-buffered saline (PBS, 1×, pH 7.4) was obtained from Thermo Fisher Scientific. Chitosan oligosaccharide lactate (average Mn ≈ 5000 Da) was purchased from Sigma-Aldrich. Cetyltrimethylammonium bromide (CTAB, 99%, molecular biology grade) was obtained from MilliporeSigma. All chemicals were used as received without further purification.

### 4.2. Nanoformulation synthesis

#### 4.2.1. Synthesis of oleate-capped NaYF_4_:Yb^3+^, Er^3+^

NaYF_4_ (Y, Yb^3+^: Er^3+^ = 57.5%:40%:2.5%) nanocrystals were synthesized by a combination of co-precipitation and thermal decomposition methods of rare-earth trifluoro-acetates in the mixture of oleic acid and 1-octadecene using a well-established protocol for the synthesis of hexagonal-phase UCNPs with some modifications.^[52]^ Briefly, 1 mmol of precursors: YCl_3_·6H_2_O (174.4 mg, 0.575 mmol), YbCl_3_·6H_2_O (155.0 mg, 0.4 mmol), and ErCl_3_·6H_2_O (9.54 mg, 0.025 mmol) were dissolved in 5 mL of methanol by sonication and subsequently added to a mixture of oleic acid and 1-octadecene (Ratio= 1:2) in a (250 mL) three-necked flask. The stirred reaction mixture was then heated to 150 °C under argon flow. After 30 min, the vacuum was applied for a further 30 min at 150 °C to remove remaining low-boiling impurities. The Rare-earth (RE) precursor-containing reaction mixture was then cooled down to room temperature (RT) under a constant argon flow. Subsequently, a methanolic solution (10 mL) containing NaOH (100 mg, 2.5 mmol) and NH_4_F (148.2 mg, 4 mmol) was added, and the resulting suspension was heated to 120 °C for 30 min to remove excess methanol. The reaction mixture was then heated to 325 °C under reflux under a gentle argon flow and kept at this temperature for 25 min. and cooled down to room temperature. The resulting UCNPs product was precipitated by the addition of ethanol, separated by centrifugation, and washed repeatedly with ethanol and cyclohexane. The obtained nanoparticles can be dispersed well in organic solvents, such as Hexane.

#### 4.2.2. Synthesis of Up-conversion coated with Mesoporous Silica Nanoparticles (UCNPs@mSiO_2_)

Porous silica coating on UCNPs was carried out according to the reverse microemulsion method, with a few modifications.^[53]^ Briefly, hydrophobic UCNPs (30 mg) were properly dispersed in Cyclohexane (15 mL), and the resulting mixture was mixed with 20 mL of water and 0.1 g (0.274 mmol) CTAB, and the mixture was then stirred vigorously at 85° C to evaporate the cyclohexane. Then, (500 µl) of Ammonium Hydroxide (NH_4_OH) was added to the above UCNPs/CTAB aqueous solution, which was then heated to 70 °C for 30 minutes. When the temperature was stable, (400 µl) of TEOS was added dropwise for 10 minutes, and the reaction mixture was stirred for 5 h at 65° C. The product was washed with ethanol several times and dried in a Vacuum oven at 70°C. The templates were removed through a speedy and effective ion exchange method. UCNPs@SiO_2_ (100 mg) with CTAB templates were added to 100 mL of ethanol containing (0.5 g) NH_4_NO_3_ under reflux for 3 h, and the final UCNPs@mSiO_2_ products after CTAB removal were separated by centrifugation, rinsed with ethanol, and dried under vacuum.

#### 4.2.3. Synthesis of amine-modified UCNPs@mSiO_2_ (UCNPs@mSiO_2_-NH_2_)

UCNPs@mSiO_2_ (100 mg) was dispersed in (10 mL) of anhydrous toluene containing (50 μL) of APTES. The suspension was agitated at 80 °C under a nitrogen atmosphere for 20 h. The amino-functionalized UCNPs@mSiO_2_ (UCNPs@mSiO_2_-NH_2_) were centrifuged, followed by washing with ethanol three times, and then dried at 60°C in a Vacuum oven.

#### 4.2.4. UCNPs@mSiO_2_-NH_2_ loaded with Photochlor (HPPH)

Amine-functionalized porous silica-coated NaYF₄: Yb^3+^/Er^3+^ (UCNPs@mSiO₂-NH₂) nanoparticles were loaded with Photochlor (HPPH), a photosensitizer, through covalent bonding using the EDC/NHS coupling reaction. Briefly, (10 mg) of UCNPs@mSiO₂-NH₂ nanoparticles were dispersed in PBS. Separately, HPPH (0.1 mM, 1 mL) was activated with a 3-fold molar excess of EDC and a similar molar amount of NHS in dimethyl sulfoxide (DMSO) to activate the carboxylic acid groups of HPPH for covalent bonding. The prepared HPPH solution was added dropwise to the UCNPs@mSiO₂-NH₂ nanoparticle suspension to achieve the desired concentration. The resulting mixture was stirred at room temperature for 24 hours in dark conditions to facilitate the coupling reaction and avoid degradation of the PS. After the reaction, the final HPPH-loaded nanoparticles were purified using a dialysis cassette (10,000 Da cutoff) to remove unreacted HPPH and any residual coupling agents, ensuring the purity of the final compound.

#### 4.2.5. Design and Synthesis of PSMA-targeted Chitosan for Targeted Delivery of Nanoparticles

In this study, PSMA-targeted chitosan was synthesized in our group based on the method described previously ^[54, 55]^, with minor modifications. This targeted chitosan was designed to enhance specificity and binding affinity for the extracellular domain of PSMA, which is overexpressed in certain treatment-resistant PCa, making it an ideal candidate for targeted drug delivery for PCa. The synthesis began with the preparation of Ot-Bu-Lys(Z)-Urea-Glu-(Ot-Bu) ₂, in which functional groups were protected to ensure controlled reactivity. The Cbz group of lysine was subsequently removed, followed by tert-butyl deprotection, affording the desired intermediate.

In parallel, azido-functionalized chitosan was prepared for the final click chemistry step.^[56, 57]^ To achieve this, N-phthaloyl chitosan (CS-Ph) was synthesized as a protective derivative of chitosan’s amino groups. This protection enabled bromination, yielding 6-bromo-6-deoxy-N-phthaloyl-chitosan (CS-Ph-Br), which was further converted into 6-azido-6-deoxy-N-phthaloyl-chitosan (CS-Ph-N₃). For the PSMA-targeted component, a click-compatible analog, DBCO-PEG₄-Urea-Glu, was synthesized. The DBCO group facilitated efficient, copper-free click conjugation with CS-Ph-N₃. Finally, the phthaloyl group was removed, yielding the PSMA-targeted chitosan product, which was purified and collected by lyophilization. Each step in this route was designed to maintain strict control over functional groups and ensure efficient conjugation between chitosan and the PSMA-targeting moieties. The use of phthaloyl and tert-butyl protecting groups prevented undesired side reactions, allowing precise modification. Collectively, these controlled steps yielded a final product tailored for PSMA receptor specificity, enhancing the potential of chitosan for targeted therapeutic delivery.

#### 4.2.6. Preparation of PSMA-targeted chitosan-capped UCNPs@mSiO_2_/HPPH

The stepwise synthesis and surface functionalization of PSMA-targeted chitosan-capped UCNPs@mSiO_2_/HPPH are given in Figure 1. Briefly, for the preparation of a 1:1 mixture of PSMA-targeted chitosan and non-targeted chitosan were dissolved in 1% acetic acid and allowed to be activated for 12 hours at room temperature. This balanced ratio was chosen to enhance cellular uptake by allowing both PSMA-specific targeting and broader electrostatic interactions with cell membranes, thus improving the overall efficiency and stability of the nanoparticles in biological environments. This mixture was then added to the UCNPs@mSiO_2_/HPPH suspension dropwise, followed by stirring at room temperature for 12 hours. Finally, the obtained chitosan-capped nanoparticles were isolated by centrifugation and washed thoroughly with water and ethanol, re-suspended in PBS, and kept at 4°C for further use.

### 4.3. Acellular evaluation and experimental design strategy

Before initiating in vitro experiments, the intrinsic PDT, SDT, catalytic, and photothermal properties of the UCNPs@mSiO₂/HPPH@TCS nanoplatform were systematically investigated under acellular conditions. These assays were intentionally performed in a cell-free environment to enable precise quantification of ^1^O₂ and ^•^OH generation, catalytic reactivity, and heat conversion efficiency following both NIR light and US activation, independent of biological variability or cellular stress responses. Establishing these fundamental physicochemical behaviors was essential to confirm functional activation of the nanoplatform and to define safe and effective exposure parameters before biological evaluation.

Acellular studies were conducted to quantify ROS production under PDT and SDT modalities across a range of exposure times. To rationally optimize treatment conditions and minimize empirical trial and error, an RSM-based experimental design was employed.^[58]^ For SDT optimization, three independent variables, US frequency, acoustic intensity, and exposure duration, were systematically evaluated using an RSM framework to predict the optimal therapeutic operating point. The experimental runs generated by the design matrix were subsequently executed in an acellular format, and the resulting ROS data were incorporated into the statistical model to identify conditions that maximized therapeutic output.

In parallel, a similar RSM approach was applied to the PDT modality using 980 nm NIR laser excitation. In this case, laser power density (0.63 W.cm^-2^) and irradiation time were selected as the two independent variables. The objective of this design was not only to maximize photodynamic activation but also to identify a safe operational window in which the photothermal effect remained within a biologically tolerable range, specifically maintaining the system temperature below 37 °C. Following experimental design generation, all PDT runs were conducted under acellular conditions, and the resulting data were analyzed using a D-optimal modeling approach to determine the best-fit predictive model and optimal exposure parameters. Collectively, this acellular, RSM-guided strategy enabled data-driven optimization of both SDT and PDT activation conditions, ensuring controlled ROS generation and thermal safety before transitioning to in vitro therapeutic assessments. Subsequent sections describe the detailed experimental procedures and analytical methods used for each modality.

#### 4.3.1. Optimization of 980 nm PDT Parameters and Photothermal Safety Assessment Using Response-Surface Methodology

To identify safe and optimal irradiation parameters for 980 nm laser-mediated PDT, RSM was employed to evaluate laser-induced photothermal effects of the nanoparticle formulation. A D-optimal experimental design was generated using Design-Expert^®^ software (Stat-Ease, Inc., Minneapolis, MN, USA), incorporating two independent variables: laser power density (categorical; 0.5 W = 0.63 W.cm^-2^ or 1W=1.27 W.cm^-2^) and irradiation time (continuous; 15-180 s). Irradiation time was treated as a continuous factor to capture nonlinear thermal accumulation behavior, while laser power was modeled as a two-level categorical variable. A quadratic polynomial model was preselected to account for curvature and interaction effects between variables. For each experimental condition, nanoparticle dispersions (3 µg. mL^-1^) were loaded into sealed quartz capillary tubes to ensure consistent optical path length and minimize convective heat loss. Samples were irradiated using a continuous-wave 980 nm laser according to the design matrix. Real-time temperature changes were monitored and recorded at predefined time intervals using an IR thermal imaging camera positioned perpendicular to the irradiation axis. The maximum temperature reached during irradiation was extracted for each condition and used as the response variable.

Photothermal behavior was evaluated under identically ambient conditions, and unirradiated samples served as negative controls to account for baseline thermal drift. All measurements were performed in triplicate, and temperature data were reported as mean ± standard deviation. The resulting dataset was analyzed using Design-Expert to generate response surface plots and predictive models. Optimal irradiation parameters were defined as those producing sufficient laser exposure for UCNPs excitation while maintaining the maximum temperature below 37 °C, thereby minimizing photothermal contributions and ensuring that observed biological effects in subsequent PDT experiments were attributable primarily to photodynamic mechanisms rather than thermal heating.

#### 4.3.2. Optimization of SDT Parameters Using RSM

To systematically optimize sonodynamic ROS generation, the RSM approach was used. A D-optimal experimental design was constructed in Design-Expert^®^ software, incorporating three factors: US frequency (categorical; 1 or 3 MHz), acoustic intensity (1-5 W/cm²), and exposure time (1-5 min). Intensity and exposure time were modeled as continuous variables, while frequency was treated as a two-level categorical factor. The final design consisted of 19 experimental runs, supplemented with five pure-error replicates and five lack-of-fit points. A quadratic model was preselected to capture curvature and interaction effects. For each design point, ROS generation was quantified using 1,3-Diphenylisobenzofuran (DPBF). A fixed concentration of DPBF was mixed with nanoparticle dispersions under identical solvent conditions. Samples were sonicated according to the design matrix, and absorbance at 410 nm was measured immediately after treatment. ROS production was calculated as the percentage decrease in DPBF absorbance relative to non-sonicated controls. All measurements were performed in triplicate, and results were reported as mean ± standard deviation. (It should be noted that DPBF predominantly reacts with singlet oxygen but may also undergo oxidation in the presence of other highly reactive oxygen species in acellular environments. Therefore, the observed DPBF degradation reflects the relative total ROS generation rather than selective ^1^O₂ quantification.

#### 4.3.3. Singlet Oxygen (¹O₂) Generation Assay

Singlet oxygen production under PDT and SDT conditions was assessed using DPBF, 30 µM as a chemical probe. Nanoparticle dispersions (3 µg. mL^-1^ HPPH equivalent) were mixed with DPBF and exposed either to NIR laser irradiation (980 nm) or US for defined durations and energy doses. The decrease in DPBF absorbance, measured by UV–Vis spectrophotometry, was used as an indicator of ¹O₂ generation. (It should be noted that selective singlet oxygen analysis was conducted separately under controlled PDT and SDT conditions, whereas the Design-Expert study aimed solely at identifying optimal operational parameters that maximize overall ROS generation).^[59, 60]^

#### 4.3.4. Peroxidase-Like (POD) Catalytic Activity

To evaluate POD-like catalytic behavior under PDT or SDT activation, methylene blue (MB) degradation was quantified. MB solution (10 µg. mL^-1^) was added to UCNPs@mSiO₂/HPPH@TCS dispersions (3 µg. mL^-1^) in the presence of H₂O₂ (1 mM). Samples were irradiated with NIR light or exposed to US for defined time intervals, and MB degradation was monitored through spectral analysis using a UV–Vis spectrophotometer.

#### 4.3.5. Acellular Tissue-Depth Penetration and ROS Generation Assay

To evaluate the penetration efficiency of 980 nm NIR light, 665 nm light, and US, an acellular ROS assay was performed using DPBF as a ROS-reactive probe. A reaction mixture containing UCNPs@mSiO₂/HPPH@TCS nanoparticles (3 µg. mL^-1^ ) and DPBF (50 μM) in ethanol was prepared in a total volume of 1 mL and transferred into the upper chamber of a 12-well Transwell insert (polycarbonate membrane, ∼10 μm thick), which allowed efficient transmission of both light and US while holding the sample in a fixed optical path. The initial DPBF absorbance at 410 nm was recorded as A₀. Fresh beef tissue slices of defined thicknesses, 1, 3, 5, 10, and 20 mm (equivalent to 0-2 cm) or none (no tissue) were placed between the irradiation source and the Transwell membrane for depth-dependent testing. For PDT experiments, the samples were irradiated either with a 980 nm continuous-wave laser or with a 665 nm laser at an identical dose of 113.4 J·cm⁻² for 3 min. For SDT experiments, samples were exposed to US at 3 MHz (5 W.cm^-2^) for 3 min using a transducer acoustically coupled to the tissue with US gel. Immediately after laser or US exposure, the DPBF absorbance at 410 nm (Aₜ) was measured using a UV–Vis spectrophotometer. ROS generation was quantified using the standard formula %ROS = [(A₀ – Aₜ) / A₀] × 100. All experiments were carried out under dark conditions to prevent unintended photodegradation of DPBF, and each condition was performed in triplicate.^[61]^

### 4.4. In Vitro Studies

#### 4.4.1. Cells and Culture

PC-3 and LNCaP (ATCC, Manassas, Virginia) PCa cell lines were selected to represent distinct biological phenotypes and PSMA expression profiles. PC-3 cells, derived from a bone metastasis of a grade IV adenocarcinoma in a 62-year-old male, are androgen-independent, highly aggressive, and PSMA-negative. By contrast, LNCaP cells originate from a left supraclavicular lymph node metastasis of a 50-year-old male and are androgen-sensitive and PSMA-positive, providing a relevant model of early-stage, hormone-dependent disease. Using both lines allowed assessment of nanoparticle targeting, uptake, and therapeutic response across biologically divergent PCa types. Cells were cultured in RPMI-1640 medium (Gibco, Invitrogen, Carlsbad, CA, USA) supplemented with 10% fetal bovine serum and 1% penicillin–streptomycin and maintained at 37 °C in a humidified 5% CO₂ incubator. For subculturing, confluent monolayers were rinsed with 1× D-PBS (ATCC 30-2200), exposed briefly to Trypsin-EDTA (ATCC 30-2101), and incubated until cell detachment. The cell suspension was neutralized with complete medium, gently triturated, and reseeded at a 1:3 split ratio. Medium was replaced two to three times weekly, and cells were used within recommended passage limits for all experiments.

#### 4.4.2. Cancer Cell Uptake

For intracellular imaging, PC-3 & LNCaP cells were seeded at a density of 5 × 10^3^ cells/well in (500 μL) of complete medium in 35 mm glass bottom cell culture dishes (polystyrene, sterile; NEST Biotechnology Co., Ltd., Wuxi, China; Cat. No. 80100, USA) and cultured for 24 h at 37 °C in 5% CO_2_. The medium was then replaced with fresh medium containing UCNPs@mSiO_2_/HPPH@TCS (0.003 mmol HPPH) and incubated for designated time intervals (0, 1, 3, 6, 12, 15, and 24 hours). Finally, the media in the chambers were replaced once again with fresh media before the cells were imaged on an Echo fluorescence microscope (Echo Laboratories, San Diego, CA) scanning microscope equipped with a 63× Plan-Apo/1.3 NA oil immersion objective with DIC capability. Further, the nucleus was counter-stained with 4,6-diamidino-2-phenylindole (DAPI) for 10 minutes and washed. Subsequently, the cells were imaged using fluorescence microscopy in the red channel to track the uptake of HPPH at different time points. The experiments were performed in triplicate.

#### 4.4.3. Sub-Cellular Localization of NPs in LNCaP Cells by Confocal Microscopy

Sub-cellular localization studies were done in confluent LNCaP cells only. For this purpose, cells were seeded on glass-bottom dishes and allowed to attach for 24 h before nanoparticle treatment. UCNPs@mSiO₂/HPPH@TCS nanoparticles were briefly bath-sonicated and added to the cells at the indicated working concentration for 1 h at 37 °C. Following incubation, cells were washed with warm HBSS and processed in two separate staining experiments, one using LumiTracker® Mito Green FM (Lumiprobe Corporation, USA) (100 nM) and the other using LumiTracker® Lyso Green (Lumiprobe Corporation, USA) (75 nM), each incubated for 25–30 min at 37 °C according to the manufacturer’s instructions. After staining, cells were washed and imaged live in phenol-red-free medium using a laser-scanning confocal microscope equipped with a 60× oil-immersion objective. Green channel organelle fluorescence (Mito Tracker or Lysotracker) was acquired using 488 nm excitation and 500-550 nm emission, while nanoparticle-associated HPPH fluorescence was detected using 633 nm excitation with emission collected at 650-720 nm. Sequential acquisition was used to prevent bleeding through. Z-stacks were collected using a 0.3-0.5 μm step size with identical settings across samples. Colocalization between nanoparticles and mitochondria or lysosomes was analyzed in ImageJ (Fiji) using Pearson’s correlation coefficient.

#### 4.4.4. Cell viability/Toxicity assays

Cell viability was assessed in both LNCaP and PC-3 cells using the Cell Counting Kit-8 (CCK-8; CK04, Dojindo Molecular Technologies, Inc., Rockville, MD, USA) according to the manufacturer’s instructions. Briefly, 10,000 cells in 100 µl of complete medium were seeded into each well of a 96-well plate. After cell attachment, PC-3 and LNCaP cells were treated with different concentrations of free HPPH and HPPH-loaded nanoparticles (NF; equivalent HPPH concentrations) at 1, 2, 3, 5, 10, and 20 µg. mL^-1^ to determine the safe dose in both cell lines. treatments were performed in triplicate. Cells were incubated for 24 h, after which (10 µl) of CCK-8 reagent was added to each well, and plates were further incubated at 37 °C for 2 h. Absorbance at 450 nm was recorded using a microplate spectrophotometer. Cell viability was calculated as a percentage of the untreated control after subtraction of background signals from blank wells.

#### 4.4.5. Photodynamic therapy (PDT) of PC-3 and LNCaP cells

Photodynamic therapy was performed on PC-3 and LNCaP cells by incubating them with (3 µg. mL^-1^) of free HPPH or NF (HPPH-equivalent) for the appropriate uptake time. After incubation, the nanoparticles or free HPPH were removed, and the cells were washed with PBS to eliminate excess or unbound HPPH. The medium was then replaced with phenol red-free medium, and the cells were exposed to light irradiation under dark conditions: 660 nm laser for free HPPH and 980 nm laser for NF, using varying exposure doses. Immediately after irradiation, the cells were returned to the incubator. Cell viability was assessed 24 hours later using the CCK-8 assay.

#### 4.4.6. Sonodynamic therapy (SDT) of PC3 and LNCaP cells

Sonodynamic therapy was performed on PC-3 and LNCaP cells by incubating them with (3 μg. mL^-1^) of free HPPH or NF (HPPH-equivalent) for the appropriate uptake time. After incubation, the nanoparticles or free HPPH were removed, and the cells were washed with PBS to eliminate excess or unbound HPPH. The medium was then replaced with phenol red-free medium, and the cells were exposed to ultrasound at 3 MHz under dark conditions, using different intensities and time points for each treatment group. Immediately after ultrasound exposure, the cells were returned to the incubator, and cell viability was assessed 24 hours later using the CCK-8 assay.

#### 4.4.7. Intracellular ROS Measurements

For ROS measurement studies, PC-3 and LNCaP cells were seeded at a density of 5 × 10³ cells per well in 500 µL of complete medium in 35 mm glass-bottom culture dishes and allowed to attach for 24 h at 37 °C in 5% CO₂. The medium was then replaced with fresh medium containing UCNPs@mSiO₂/HPPH@TCS (equivalent to 3 µg. mL^-1^ HPPH), and cells were incubated for the designated uptake time for each cell line at pH 7.4. Following uptake, the medium was replaced to remove extracellular nanoparticles, and cells were either exposed to NIR laser irradiation (980 nm; 113.4 j.cm^-^²) or US (3 MHz; 5.0 W, 3 min) or maintained without irradiation as a control. After treatment, the medium was replaced with fresh medium containing 5 µM 2′,7′-Dichlorodihydrofluorescein diacetate (DCFH-DA), and cells were incubated for an additional 30 min to allow intracellular probe conversion. Finally, cells were imaged using an Echo fluorescence microscope (Echo Laboratories, San Diego, CA, USA), and fluorescence intensity was quantified using Fiji software.

#### 4.4.8. Intracellular Singlet Oxygen Detection

LNCaP cells were seeded at a density of 5 × 10³ cells per well in 500 µL of complete medium in 35 mm glass-bottom culture dishes and allowed to attach for 24 h at 37 °C in 5% CO₂. The medium was then replaced with fresh medium containing UCNPs@mSiO₂/HPPH@TCS (equivalent to 3 µg. mL^-1^ HPPH), and cells were incubated for the designated uptake time for each cell line at pH 7.4. Following uptake, the medium was replaced to remove extracellular nanoparticles, and cells were either exposed to NIR laser irradiation (980 nm; 113.4 j.cm^-^²) or US (3 MHz; 5.0 W.cm^-2^, 3 min) or maintained without irradiation as a control. After treatment, the medium was replaced with fresh medium containing 5 µM SOSG (Singlet Oxygen Sensor Green), and cells were incubated for an additional 30 min to allow intracellular probe conversion. Finally, cells were imaged using an Echo fluorescence microscope (Echo Laboratories, San Diego, CA, USA), and fluorescence intensity was quantified using Fiji software.

#### 4.4.9. Gene Expression analysis

To evaluate treatment-induced transcriptional changes in inflammatory and apoptosis-related signaling pathways following NF exposure and photodynamic or sonodynamic activation, gene expression analysis was performed using quantitative real-time reverse transcription polymerase chain reaction (qRT-PCR) in LNCaP cells following appropriate treatment conditions. For this, LNCaP cells were seeded in 6-well plates at a density of 1 × 10⁵ cells/mL and allowed to adhere for 24 h at 37 °C in a humidified incubator containing 5% CO₂. Following attachment, cells were treated with NF and assigned to one of three experimental conditions: (i) NF alone without external activation (dark control), (ii) NF followed by PDT, and (iii) NF followed by SDT. After treatment and/or activation, cells were incubated for an additional 24 h prior to RNA isolation. This was followed by total RNA extraction using the acid guanidinium thiocyanate-phenol-chloroform method with TRIzol^®^ reagent according to the manufacturer’s protocol. Briefly, (1 mL) of TRIzol^®^ reagent was added to each well containing approximately 1 × 10⁵ cells, and the resulting lysates were transferred to 1.5 mL microcentrifuge tubes. Chloroform (200 μL) was added, and samples were incubated at room temperature for 10 min, followed by centrifugation at 13,000 rpm for 15 min at 4 °C. The aqueous phase was carefully collected, and RNA was precipitated with 600 μL of isopropanol. RNA samples were stored at −80 °C overnight before reverse transcription. For cDNA synthesis, 500 ng of total RNA was reverse transcribed using the All-in-One Universal RT Master Mix synthesis kit (Lamba Biotech, St. Louis, MO, USA) following the manufacturer’s instructions. qRT-PCR was performed using 1 μL of cDNA template and gene-specific primers (IL-1β, TNF-α, NF-κB, caspase-3, Bax, and BCL-2; Integrated DNA Technologies, San Diego, CA). Amplification reactions were conducted using SYBR^®^ Green master mix containing dNTPs, MgCl₂, and DNA polymerase (Bio-Rad, Hercules, CA), with a final primer concentration of 0.1 μM. Thermal cycling conditions consisted of an initial denaturation at 95 °C for 3 min, followed by 40 cycles of 95 °C for 40 s, 60 °C for 30 s, and 72 °C for 1 min, with a final extension at 72 °C for 5 min. Relative gene expression levels were calculated using the comparative threshold cycle (Ct) method. β-Actin served as the endogenous reference gene to normalize for variations in RNA input. ΔCt values were calculated as the difference between target gene Ct and β-actin Ct, and fold changes in gene expression relative to control samples were determined using the 2⁻ΔΔCt method and expressed as the transcript accumulation index (TAI).

#### 4.4.10. Spheroid Culture via Hanging-Drop Technique

3D multicellular spheroids were generated using the hanging-drop method to mimic the architectural complexity of the prostate tumor microenvironment.^[62]^ LNCaP cells were suspended in a specialized medium and dispensed as drop-casts onto the lids of small culture dishes. Upon inverting the lids, the droplets were maintained over a reservoir of deionized water to prevent evaporation and maintain local humidity within the incubator. To evaluate size-dependent therapeutic efficacy, two distinct batches were cultured to reach maturation at 6 days (small spheroids) and 14 days (large spheroids).

Mature spheroids were carefully transferred to glass-bottom dishes for high-resolution imaging and treatment. The cultures were incubated with the targeted NF (UCNPs@mSiO_2_/HPPH@TCS) in phenol red-free medium for a predetermined uptake duration of 3 hours. Subsequently, PDT and SDT were performed independently to assess the ablation efficiency of the HPPH-loaded platform in 3D structures.

Immediately, before fluorescence imaging, the spheroids were co-incubated with pSIVA-IANBD reagent (Novus Biologicals, Littleton, CO, USA). This staining approach allowed for the simultaneous detection of externalized phosphatidylserine (an early-stage apoptosis marker). A control group was maintained without the pSIVA-IANBD marker to establish baseline fluorescence. Time-lapse imaging was conducted at 0, 1, 3, 6, 12, 24, 36, and 48 h post-treatment to monitor real-time morphological changes, volumetric contraction, and the kinetics of pre-apoptotic signal intensity. To validate the structural integrity and cellular distribution within the 3D model, a separate batch of spheroids was fixed and stained with DAPI. Multi-channel fluorescence imaging was performed using the following parameters: Blue Channel (DAPI): λ_ex_ = 405 nm for nuclei visualization and confirmation of spheroid formation. Green Channel (pSIVA-IANBD): λ_ex_= 488 nm for localized pre-apoptotic signaling. Red Channel (HPPH): λ_ex_= 561nm for tracking the penetration and localization of the photosensitizer-loaded NF.

## Acknowledgements

This work was supported by the Office of the Vice President for Research and Economic Development at the University at Buffalo (Award No. 1180592-2-75023) through the Institute for Lasers, Photonics and Biophotonics, and in part by Merit Award BX005318 from the Biomedical Laboratory Research and Development (BLRD) Service of the U.S. Department of Veterans Affairs (to S.A.S. and R.A.). We gratefully acknowledge Photolitec, LLC (Buffalo, NY) for providing the HPPH photosensitizer.

## Data Availability Statement

The data that support the findings of this study are available from the corresponding author upon reasonable request.

