## Supplementary Information for "Comparative Study of Photodynamic and Sonodynamic Therapy Using a PSMA-Targeted Up-conversion Nanoplatform for Castration-Resistant Prostate Cancer"

**Table of Contents**

**Figures**

**Figure S1.** High-resolution transmission electron microscopy (HRTEM) image of the synthesized UCNPs.

**Figure S2.** FTIR spectra of nanoparticles at different stages of mesoporous silica coating before and after CTAB template removal.

**Figure S3.** DLS size distribution and zeta potential measurements of nanoparticles after mesoporous silica coating.

**Figure S4.** FTIR spectra of nanoparticles after amine functionalization, confirming successful surface modification.

**Figure S5.** Dynamic light scattering (DLS) size distribution and zeta potential analysis of the nanoparticles after amine functionalization.

**Figure S6.** Physicochemical characterization of HPPH-loaded nanoparticles by DLS and zeta potential analysis.

**Figure S7.** DLS size analysis of final nanoparticles after coating with PSMA-targeted Chitosan.

**Figure S8.** (a) Linear correlation plot predicted versus actual temperature values for the 980 nm laser-mediated PDT model. (b) Interaction plot illustrating the synergistic effects of irradiation time and laser power density on thermal elevation.

**Figure S9.** Optimization profile identifying the experimental window (165.7 s, 0.5 W) required to maintain temperature within the physiological safety threshold (~32 °C).

**Figure S10.** (a) Correlation plot for the Response Surface Methodology (RSM) model of ROS production under SDT. (b) 3D response surface plot demonstrating the multivariate interactions between sonication time, ultrasound intensity, and frequency on ROS yield.

**Figure S11.** Optimized SDT parameters (3 MHz, 5 W.cm-2, 2 min) yielded a predicted relative ROS production of 143.7%.

**Figure S12.** Evaluation of acellular singlet oxygen (^1^O₂) generation kinetics by monitoring time-dependent UV–Vis absorption spectra of DPBF in the presence of the NF under PDT (980 nm) and SDT (3 MHz, 5 W).

**Figure S13.** Assessment of Photodynamically and Sono-dynamically Generated Hydroxyl Radicals (^•^OH) via UV–Vis Absorbance Measurements.

**Figure S14.** Beef tissue phantoms with increasing thicknesses (0.1–2 cm) were placed between the irradiation/ultrasound source and the reaction system to evaluate penetration-dependent ROS generation.

**Figure S15.** Schematic and photographic representation of the acellular tissue depth penetration experimental setup used to compare photodynamic therapy at 980 nm and 665 nm, and sonodynamic therapy (3 MHz, 5 W).

**FigureS16.** Intracellular reactive oxygen species generation induced by UCNPs@mSiO₂/HPPH@TCS nanoparticles under PDT and SDT in LNCaP cells.

**FigureS17.** Intracellular reactive oxygen species generation induced by UCNPs@mSiO₂/HPPH@TCS nanoparticles under PDT and SDT in PC-3 cells.

**Tables**

**Table S1.** Analysis of variance (ANOVA) for the reduced quartic model describing the effect of irradiation time (A) and laser intensity (B) on temperature elevation during acellular photodynamic therapy (PDT) experiments.

**Table S2.** Analysis of variance (ANOVA) for the reduced quartic model evaluating the effects of ultrasound time (A), intensity (B), and frequency (C) on total reactive oxygen species (ROS) generation during acellular sonodynamic therapy (SDT) experiments.

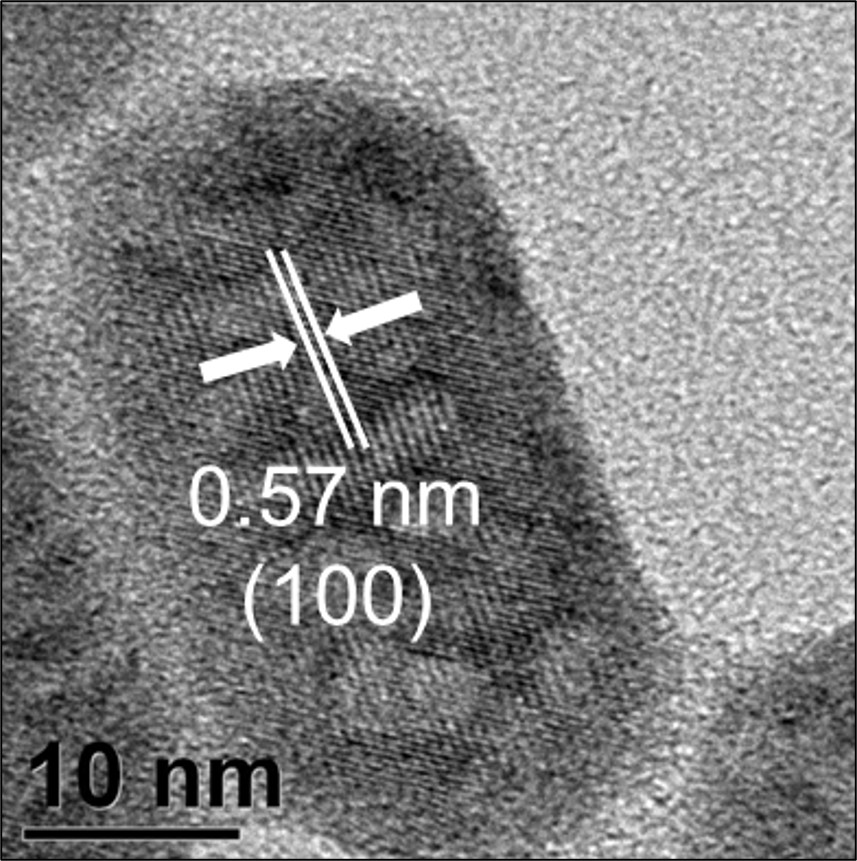

**Figure S1.** High-resolution transmission electron microscopy (HRTEM) image of the synthesized UCNPs. The lattice fringes with a d-spacing of 0.57 nm correspond to the (100) crystal plane of the hexagonal phase (e.g., β-NaYF₄), confirming the high crystallinity of the nanoparticles. The scale bar is 10 nm.

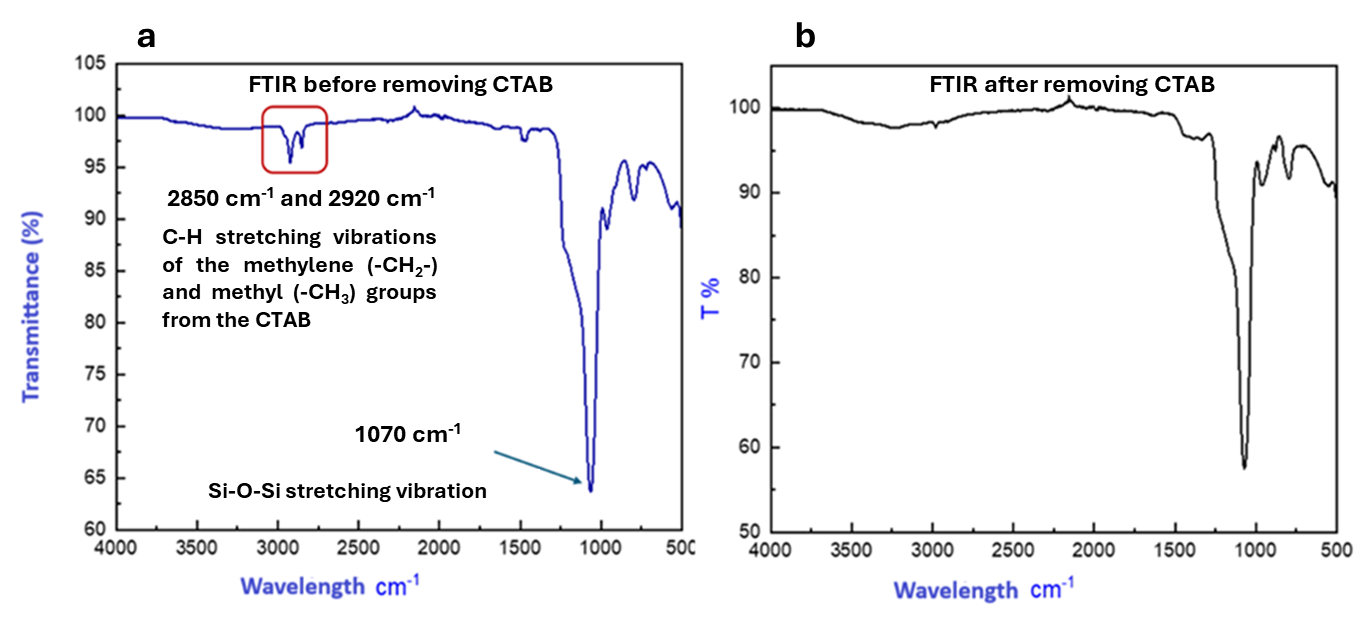

**Figure S2.** (a) FTIR spectra of nanoparticles after mesoporous silica coating, before and (b) after CTAB template removal, confirming successful surface modification at each stage.

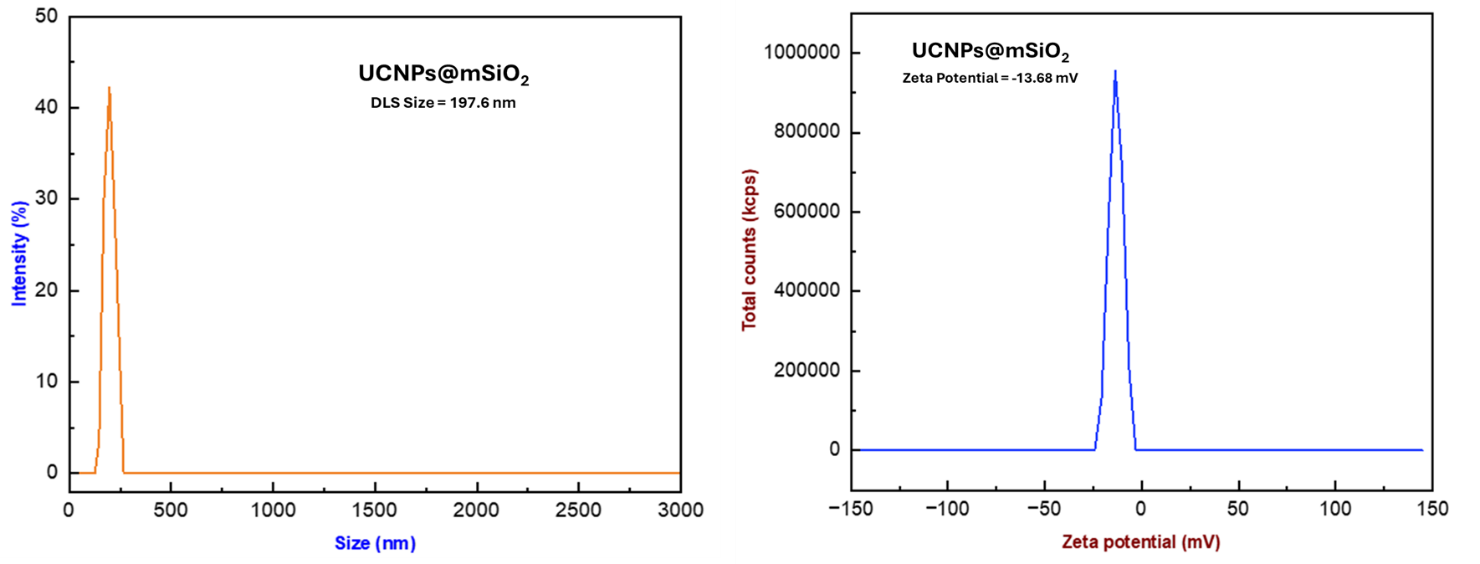

**Figure S3.** DLS size distribution and zeta potential measurements of nanoparticles after mesoporous silica coating.

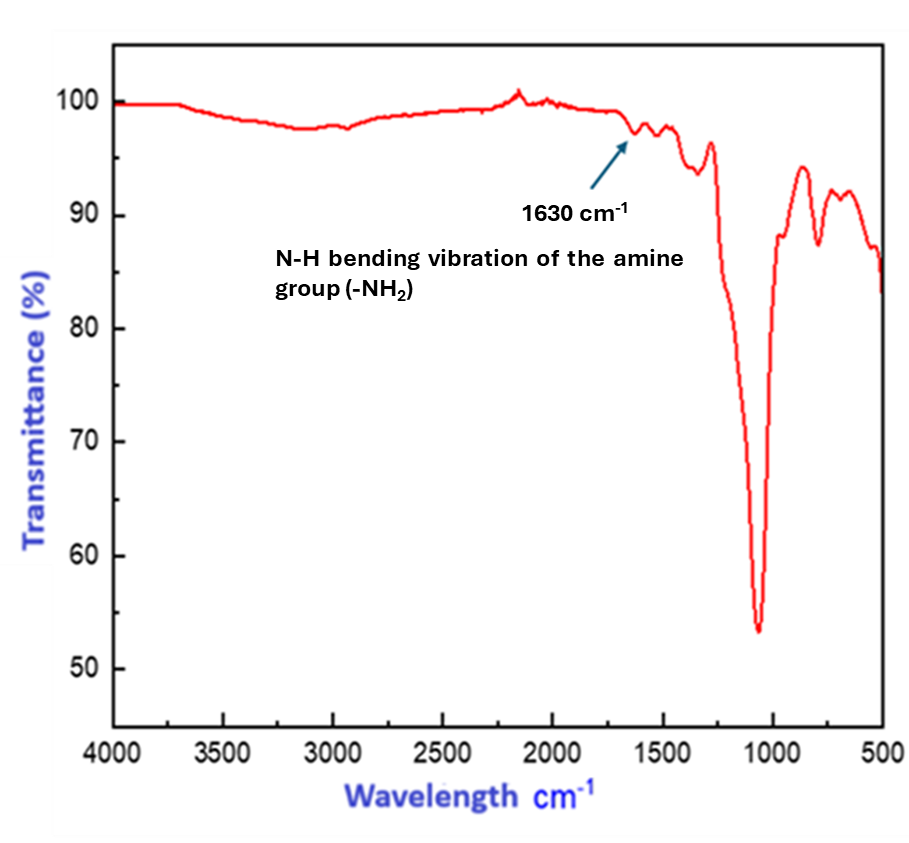

**Figure S4.** FTIR spectra of nanoparticles after amine functionalization, confirming successful surface modification.

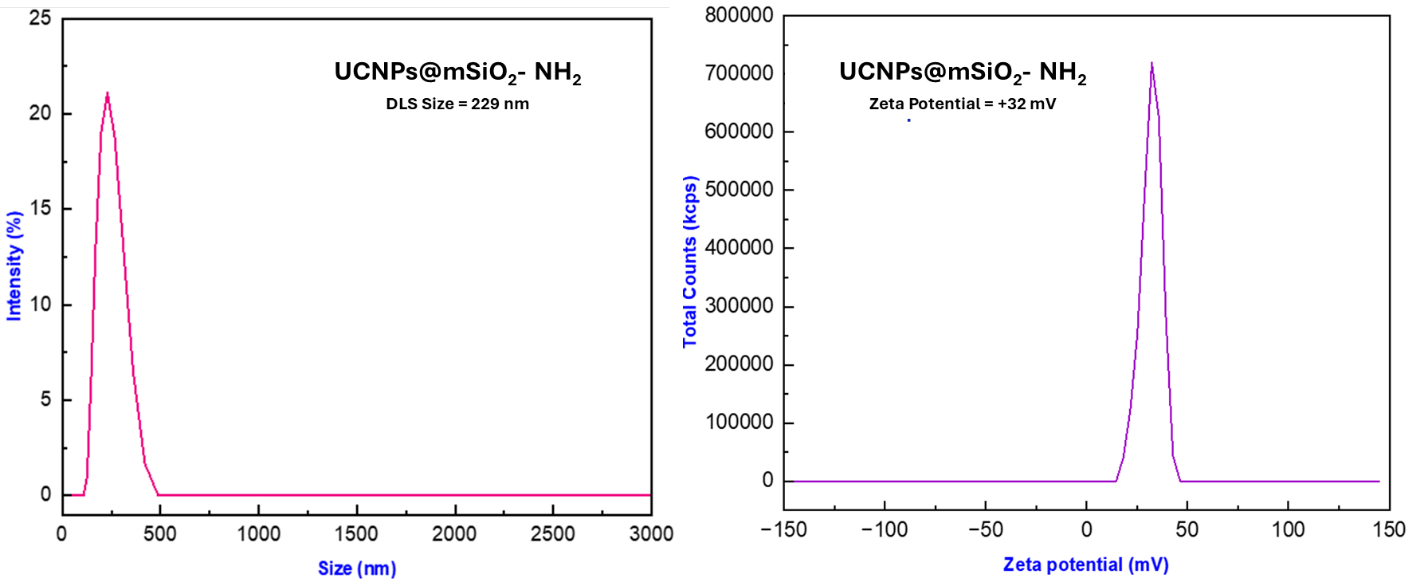

**Figure S5.** Dynamic light scattering (DLS) size distribution and zeta potential analysis of the nanoparticles after amine functionalization. The hydrodynamic diameter increased to approximately 229 nm, while the surface charge shifted to a positive value (+32 mV), confirming successful introduction of surface amine (–NH₂) groups.

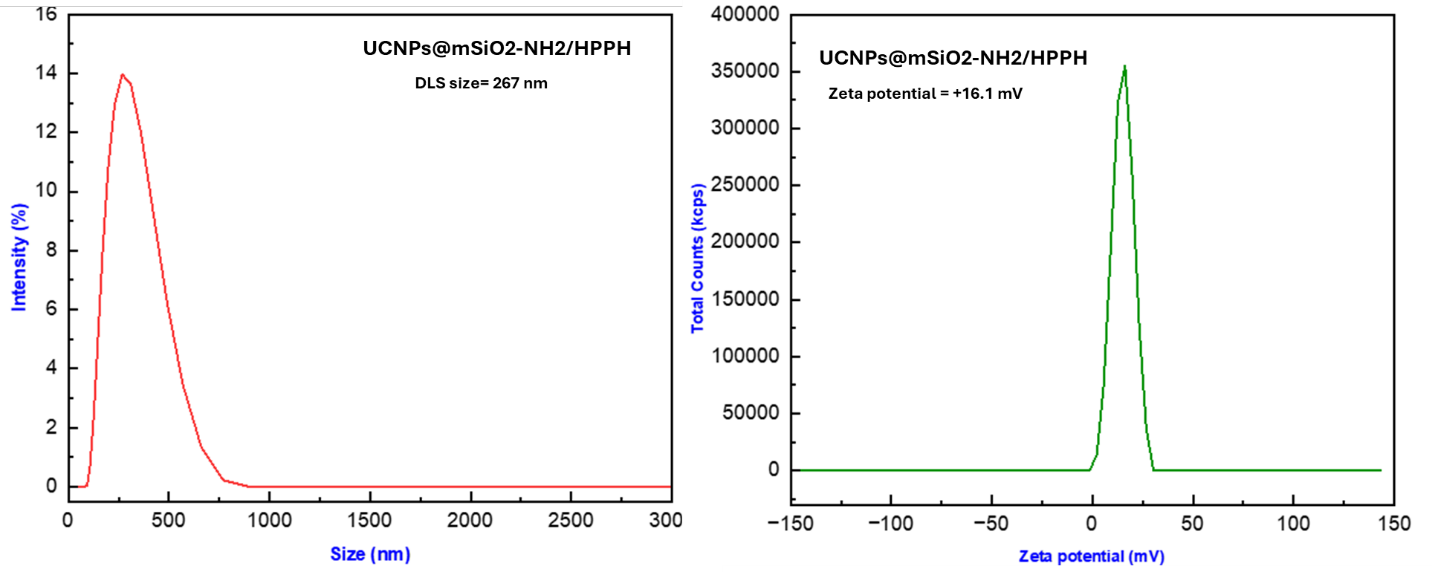

**(b)**

**(a)**

**Figure S6.** Physicochemical characterization of HPPH-loaded nanoparticles by DLS and zeta potential analysis. (a) DLS and (b) zeta potential characterization of HPPH-loaded nanoparticles showing increased hydrodynamic size (~267 nm) and reduced positive surface charge (+16.1 mV) due to interaction between surface amines and HPPH carboxyl groups.

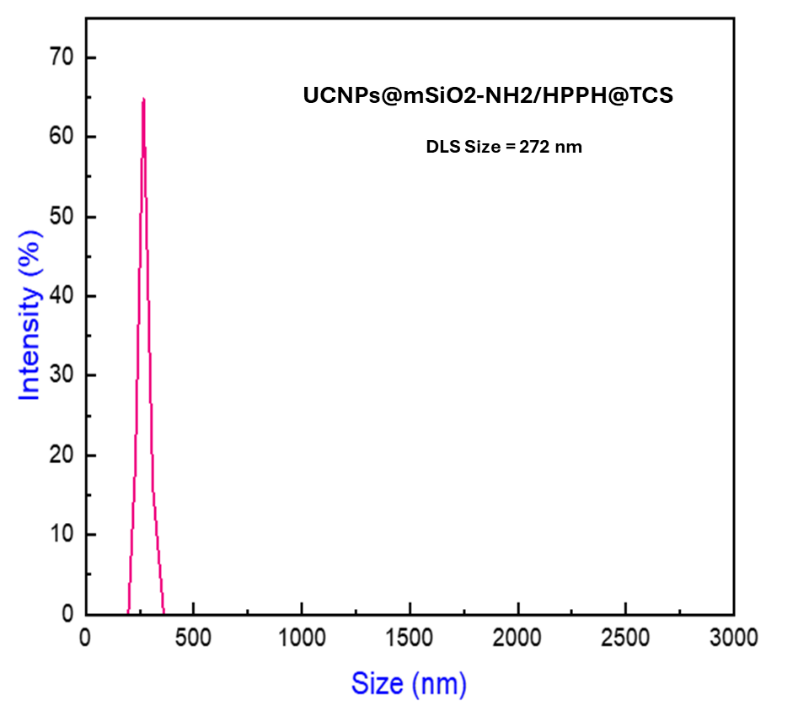

**Figure S7.** DLS size analysis of final nanoparticles after coating with PSMA-targeted Chitosan.

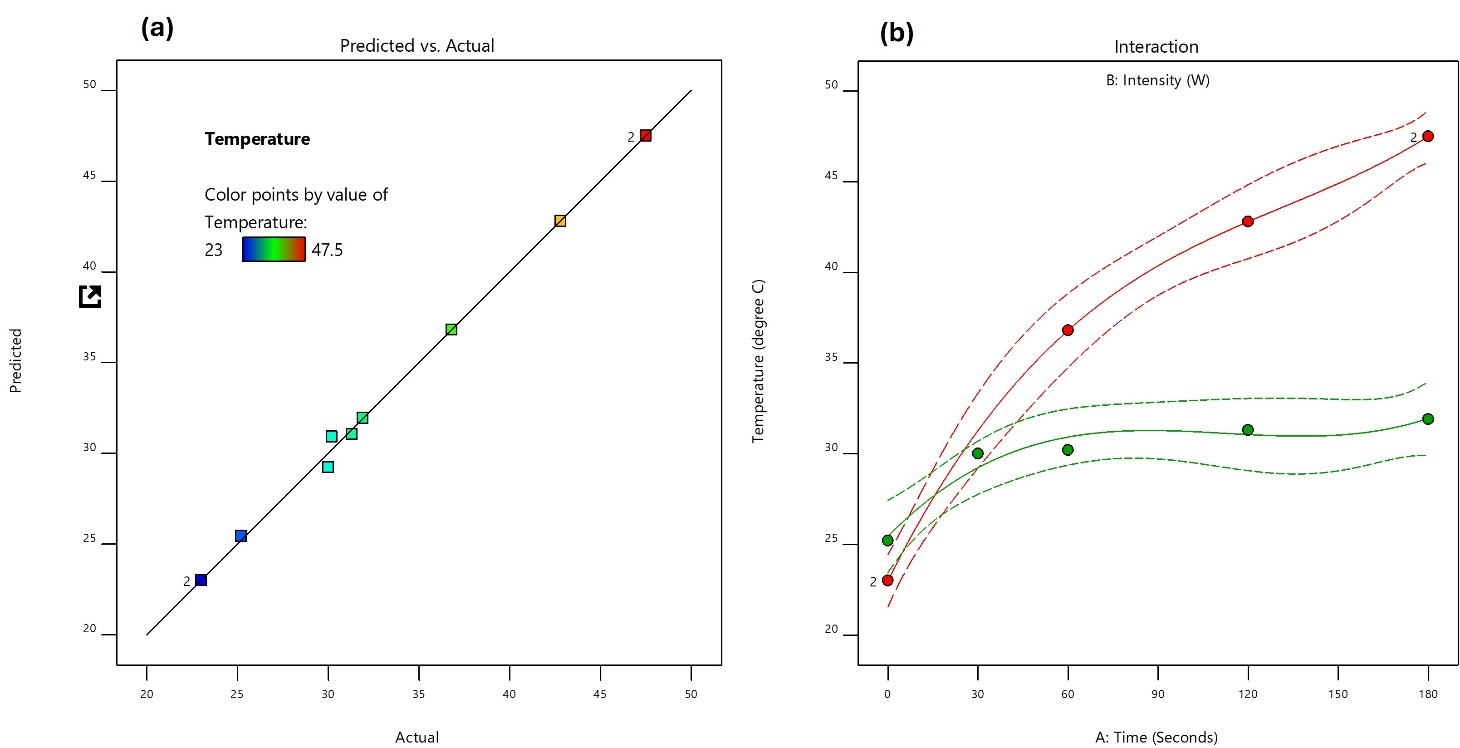

**Figure S8.** (a) Linear correlation plot predicted versus actual temperature values for the 980 nm laser-mediated PDT model. (b) Interaction plot illustrating the synergistic effects of irradiation time and laser power density on thermal elevation.

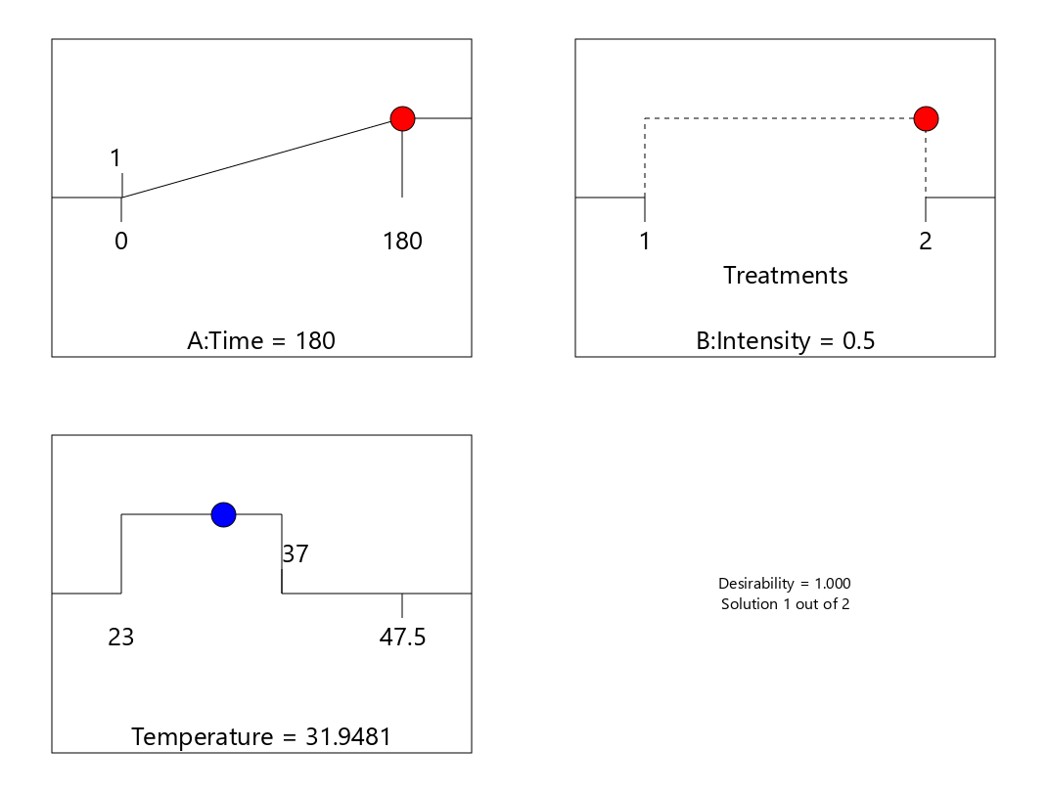

**Figure S9.** Optimization profile identifying the experimental window (165.7 s, 0.5 W) required to maintain temperature within the physiological safety threshold (~32 °C).

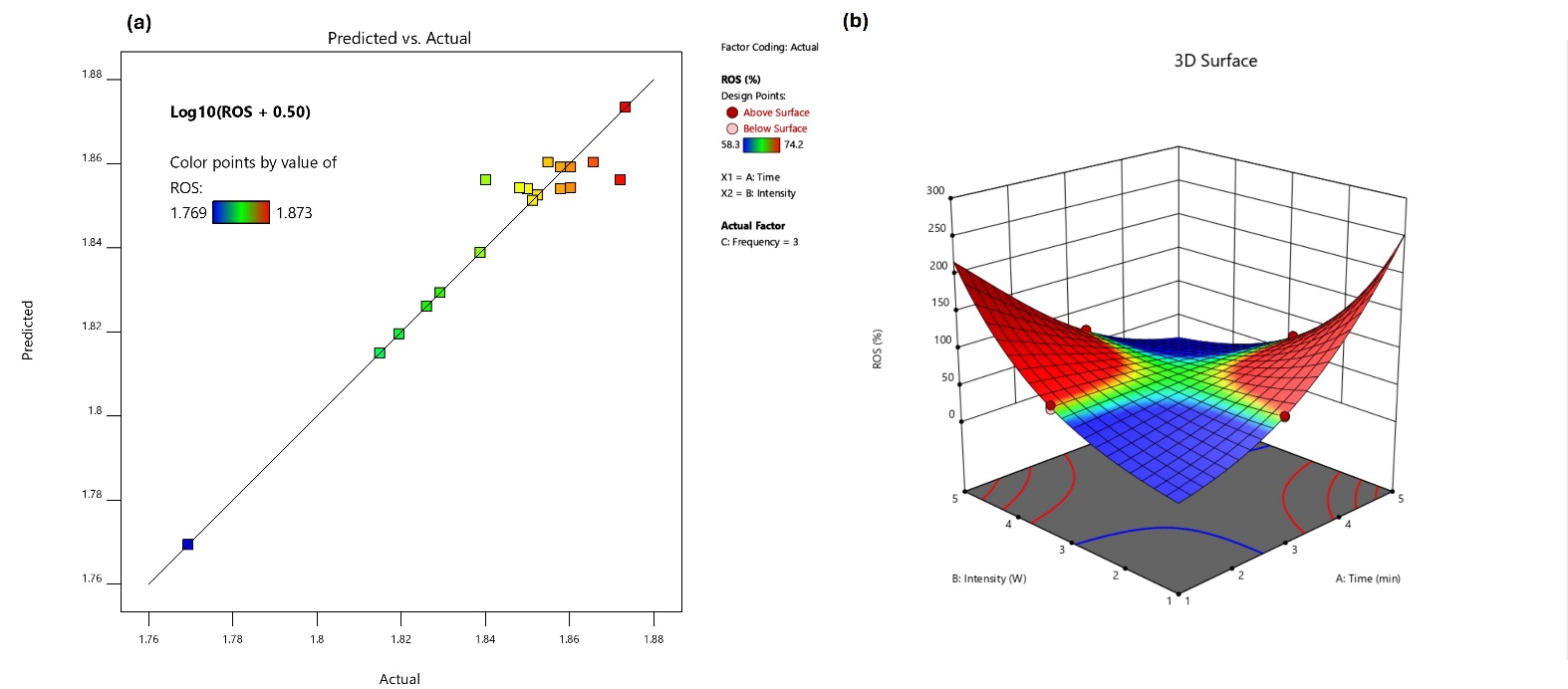

**Figure S10.** (a) Correlation plot for the Response Surface Methodology (RSM) model of ROS production under SDT. (b) 3D response surface plot demonstrating the multivariate interactions between sonication time, ultrasound intensity, and frequency on ROS yield.

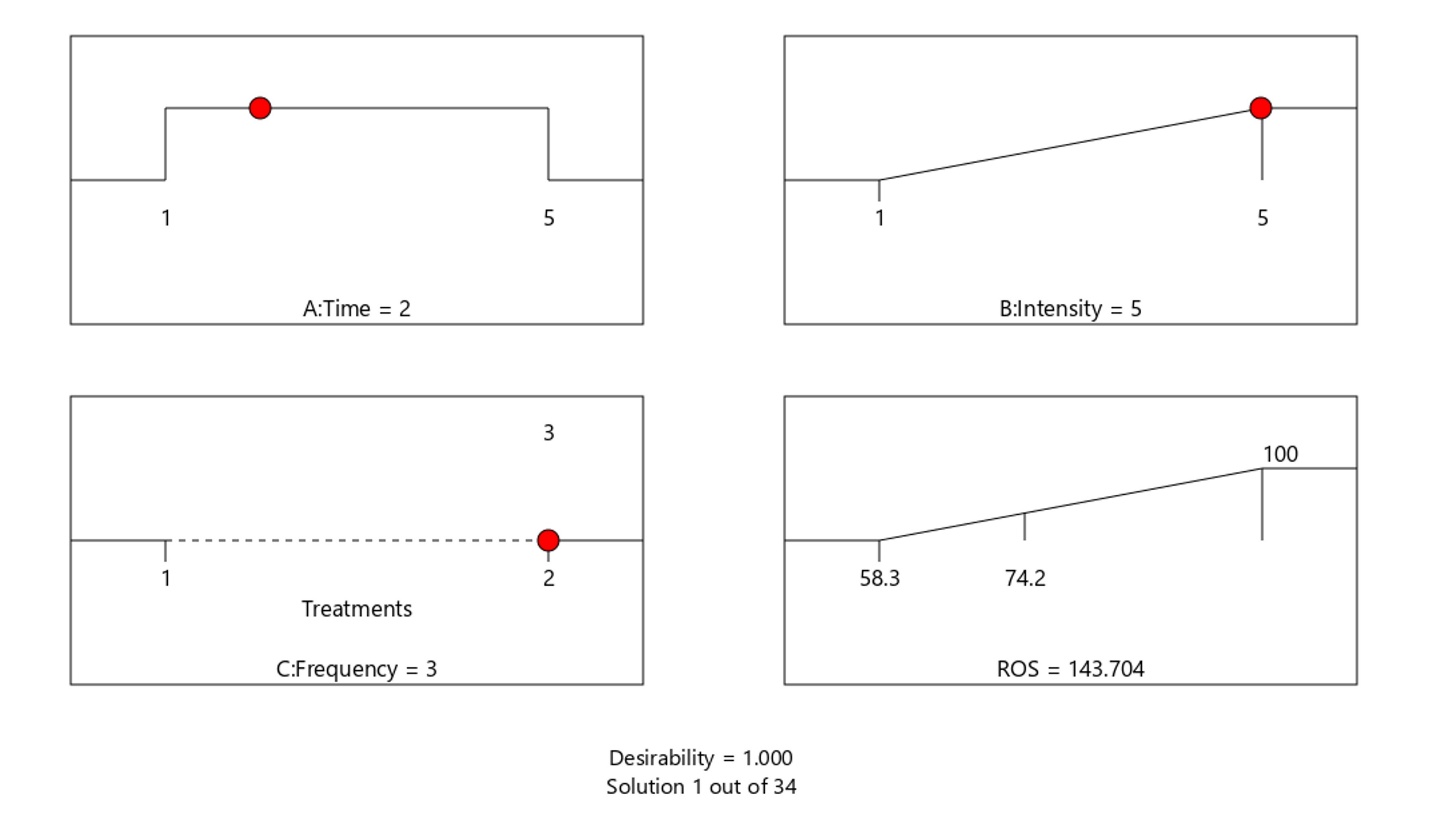

**Figure S11.** Optimized SDT parameters (3 MHz, 5 W.cm-2, 2 min) yielded a predicted relative ROS production of 143.7%.

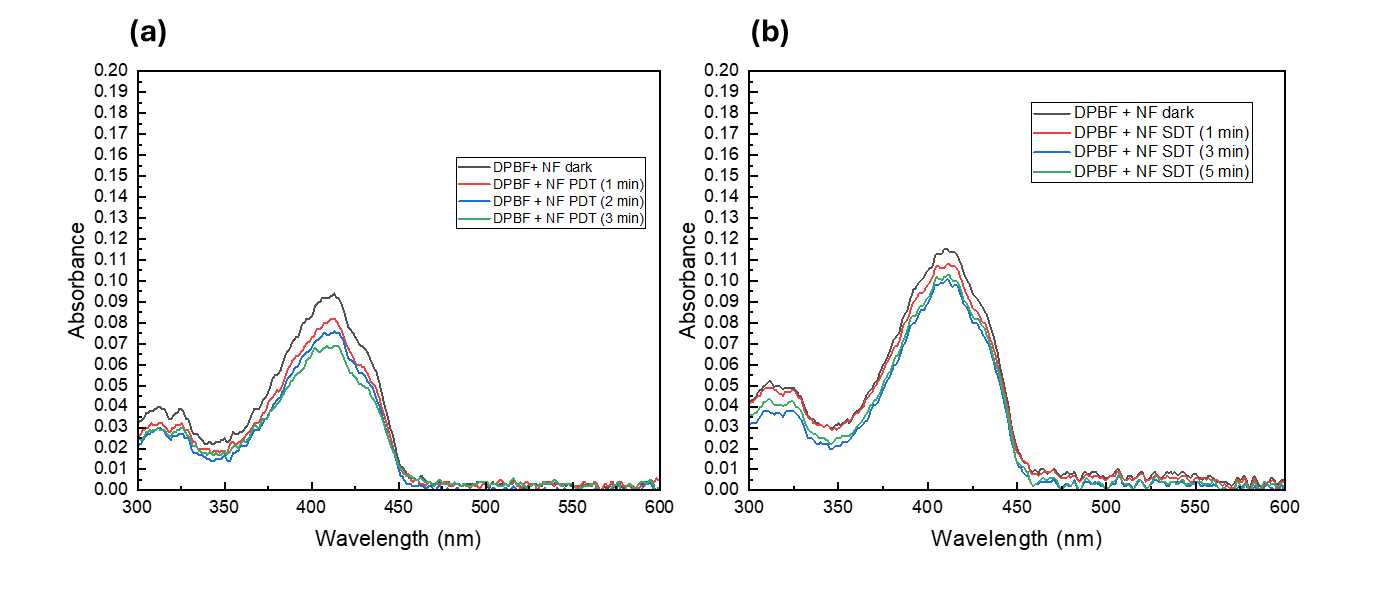

**Figure S12.** Evaluation of acellular singlet oxygen (^1^O₂) generation kinetics by monitoring time-dependent UV–Vis absorption spectra of DPBF in the presence of the NF under PDT (980 nm) and SDT (3 MHz, 5 W). (a) The systematic and time-resolved decrease in the characteristic DPBF absorbance peak at λ= 410 nm confirms the successful energy transfer from the UCNP core to the HPPH photosensitizer and subsequent ^1^O_2_ production. (b) Time-resolved UV–Vis absorption spectra of DPBF in the presence of the NF under ultrasound (US) irradiation (3 MHz, 5 W). The progressive attenuation of the characteristic absorbance peak at λ= 410 nm confirms the sonodynamically induced generation of ^1^O_2_.

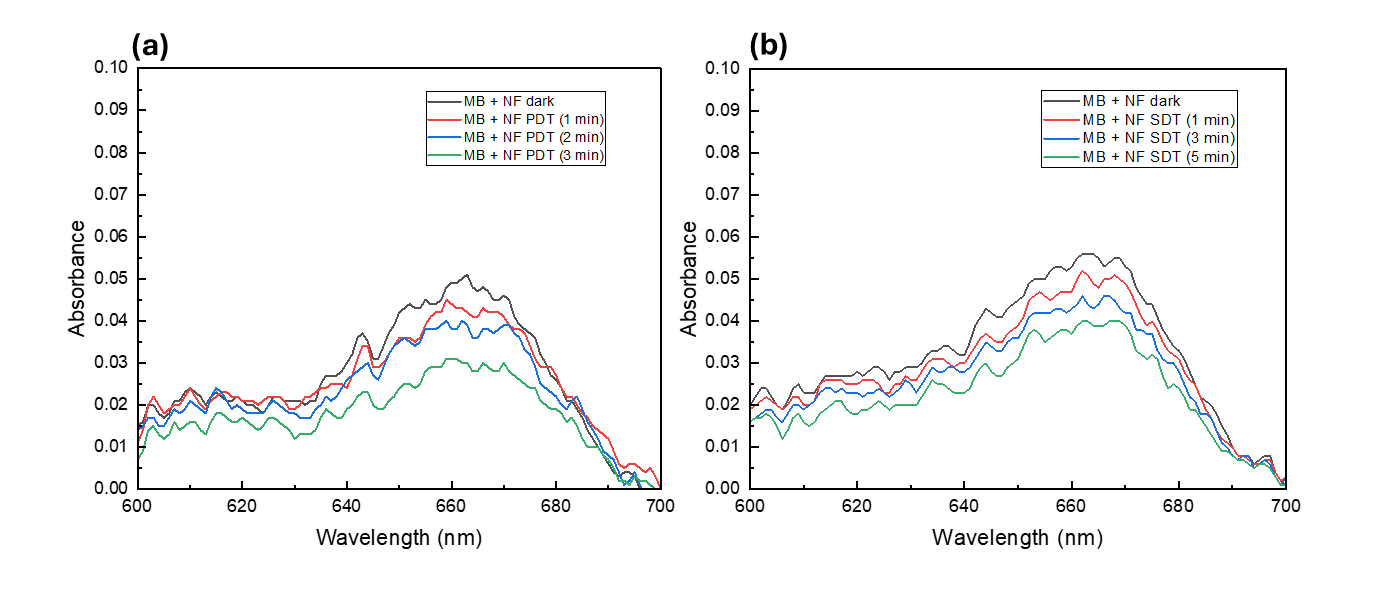

**Figure S13.** Assessment of Photodynamically and Sonodynamically Generated Hydroxyl Radicals (^•^OH) via UV–Vis Absorbance Measurements. (a) Time-dependent UV–Vis absorption spectra of Methylene Blue (MB) in the presence of the NF under 980 nm NIR laser irradiation. The systematic decrease in the characteristic MB absorbance peak at λ= 664 nm serves as an indicator of ^•^OH-mediated oxidative degradation. (b) Time-resolved UV–Vis absorption spectra of MB incubated with the NF during ultrasound (US) exposure (3 MHz, 5 W). The progressive attenuation of the 664 nm peak over a 5-minute sonication period reflects the robust generation of ^•^OH species via the sonocatalytic activation of the NF.

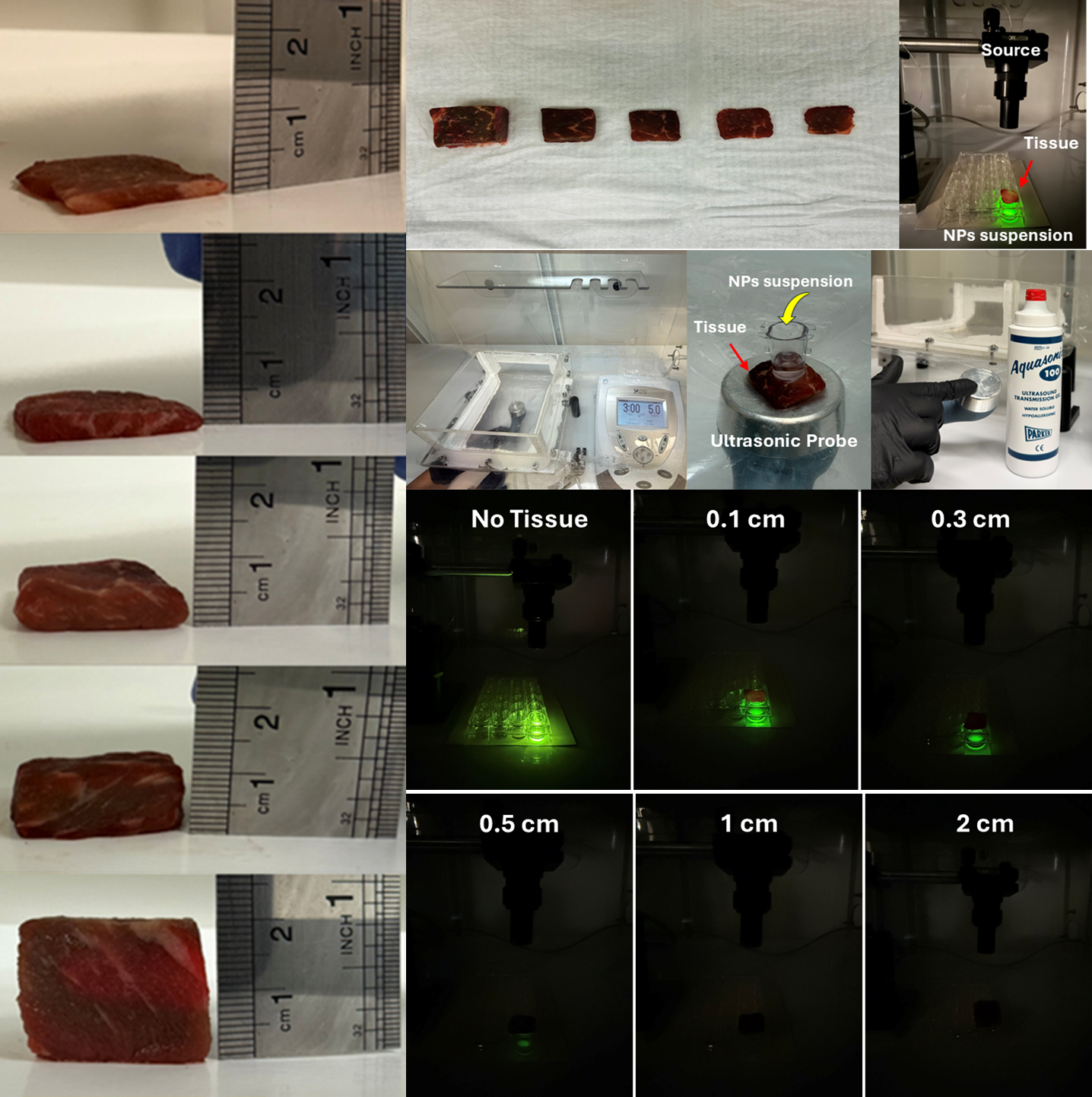

**Figure S14.** Beef tissue phantoms with increasing thicknesses (0.1–2 cm) were placed between the irradiation/ultrasound source and the reaction system to evaluate penetration-dependent ROS generation.

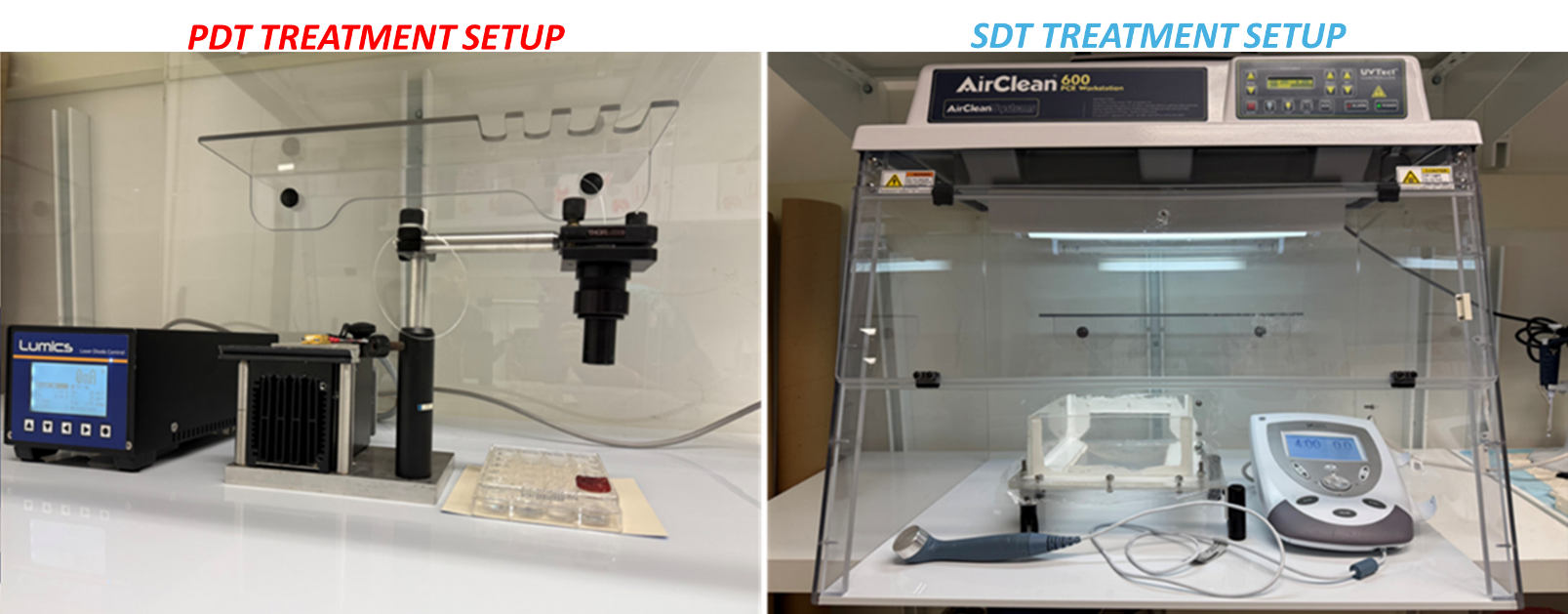

**Figure S15**. Photographic representation of the experimental setup and equipment used for photodynamic therapy (980 nm and 665 nm lasers) and sonodynamic therapy (3 MHz ultrasound) in both cellular and acellular experiments.

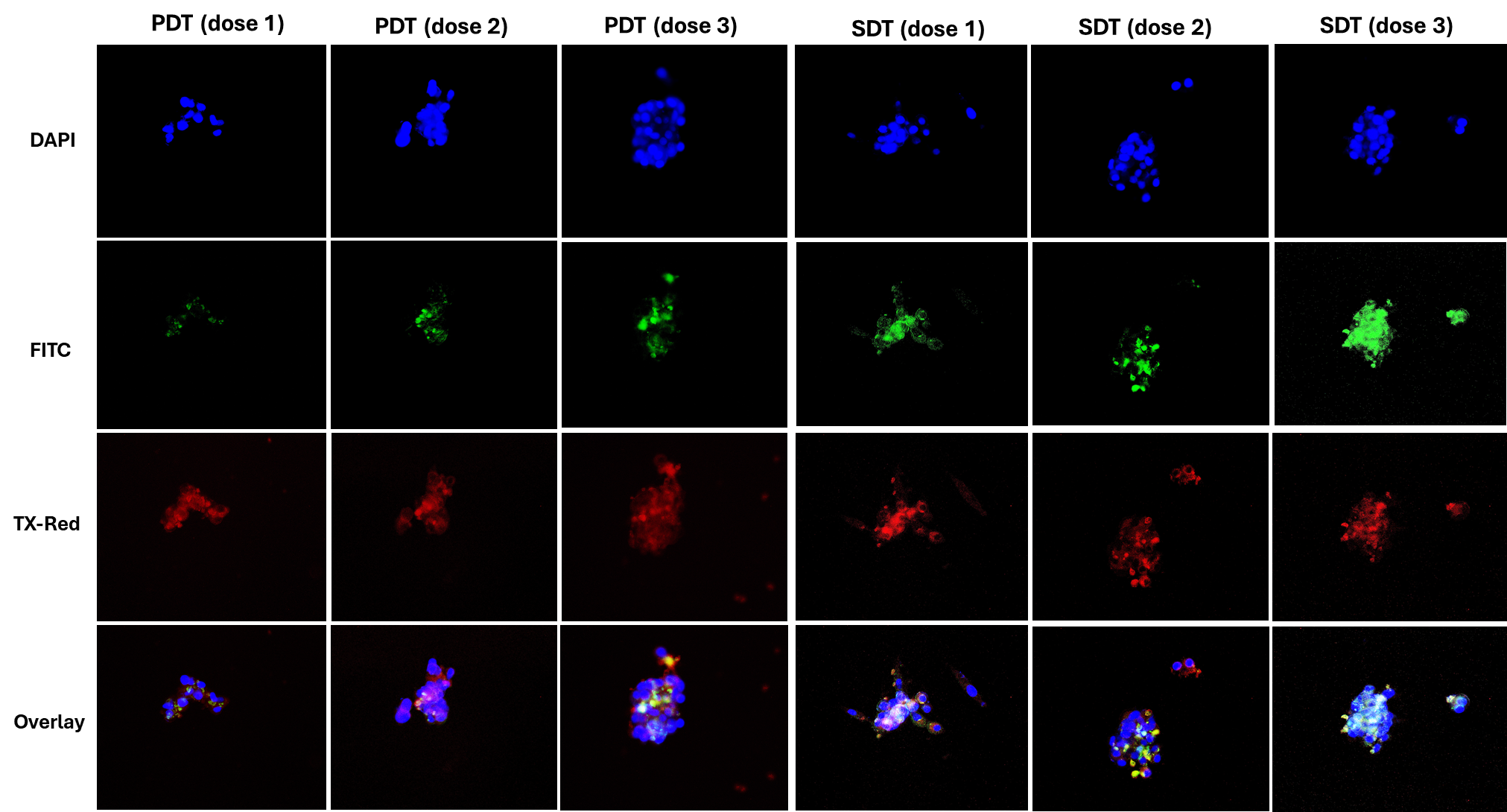

**Figure S16.** Intracellular reactive oxygen species generation induced by UCNPs@mSiO₂/HPPH@TCS nanoparticles under PDT and SDT in LNCaP cells. Cells were incubated with the NF and treated with PDT at fluences of 37.8, 75.6, and 113.4 J.cm⁻² or SDT for 1, 3, and 5 min at a fixed ultrasound frequency (3 MHz) and power (5 W). Intracellular ROS production was visualized using DCFH-DA, which produces green fluorescence upon reaction with ROS. Fluorescence channels are shown as follows: top panel (blue) – DAPI for nuclear staining, second panel (green) – DCF fluorescence for ROS detected via the FITC channel, third panel (red) – TxRed for nanoparticle-associated HPPH, and bottom panel, merged overlay of all channels. Representative fluorescence microscopy images demonstrate dose- and time-dependent ROS generation under both PDT and SDT conditions in PSMA^+^ LNCaP cells.

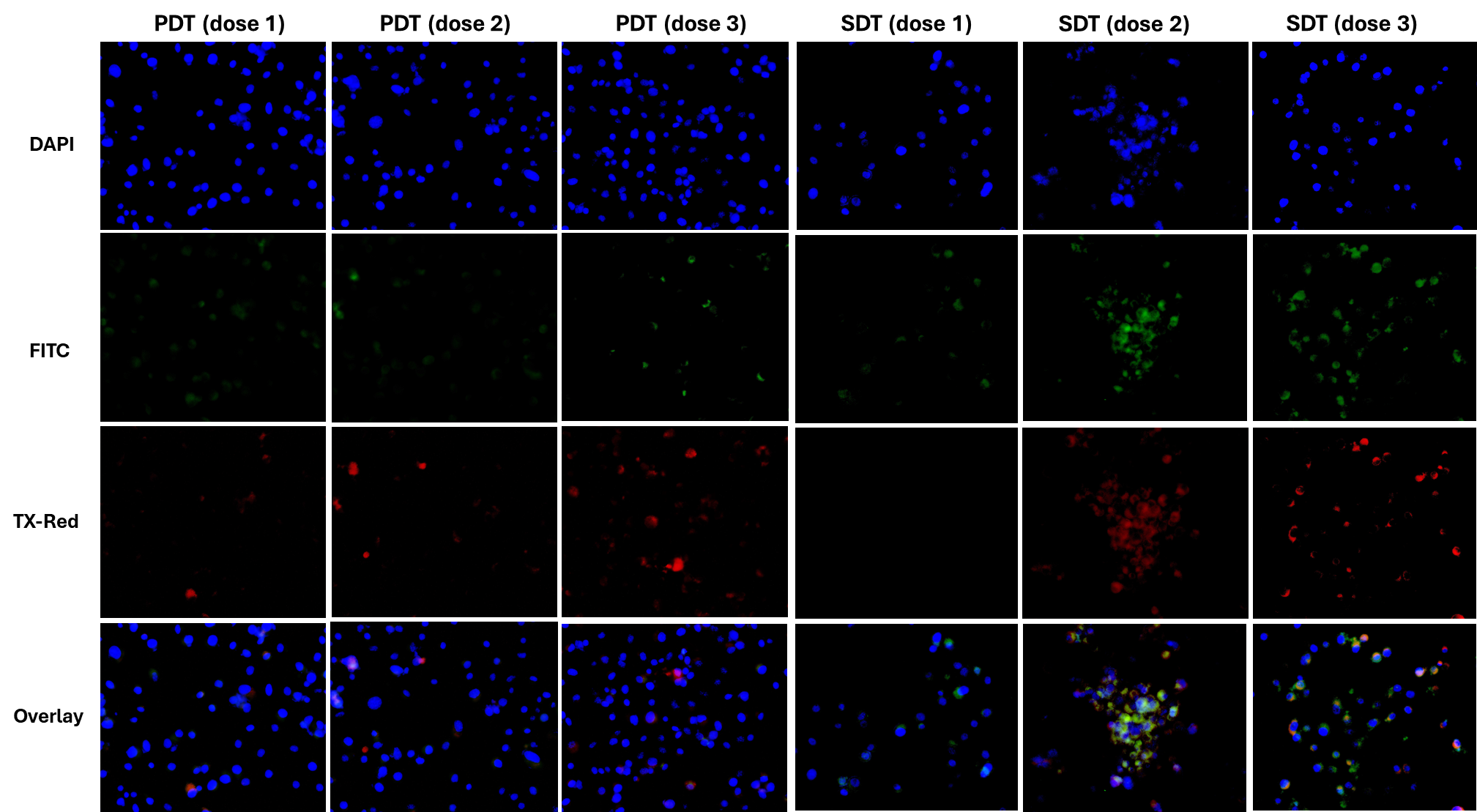

**Figure S17.** Intracellular generation of reactive oxygen species (ROS) in PC-3 cells following treatment with UCNPs@mSiO₂/HPPH@TCS (NF) under photodynamic therapy (PDT) and sonodynamic therapy (SDT) conditions. Cells were incubated with NF and subsequently subjected to PDT at light doses of 37.8, 75.6, and 113.4 J cm⁻² or SDT exposure for 1, 3, and 5 min using a constant ultrasound frequency (3 MHz) and power (5 W). Intracellular ROS levels were detected using DCFH-DA, which emits green.

**Table S1**. Analysis of variance (ANOVA) for the reduced quartic model describing the effect of irradiation time (A) and laser intensity (B) on temperature elevation during acellular photodynamic therapy (PDT) experiments. The model demonstrates high statistical significance (P < 0.001) with excellent goodness-of-fit parameters (R² = 0.9985, Adeq Precision = 44.83).

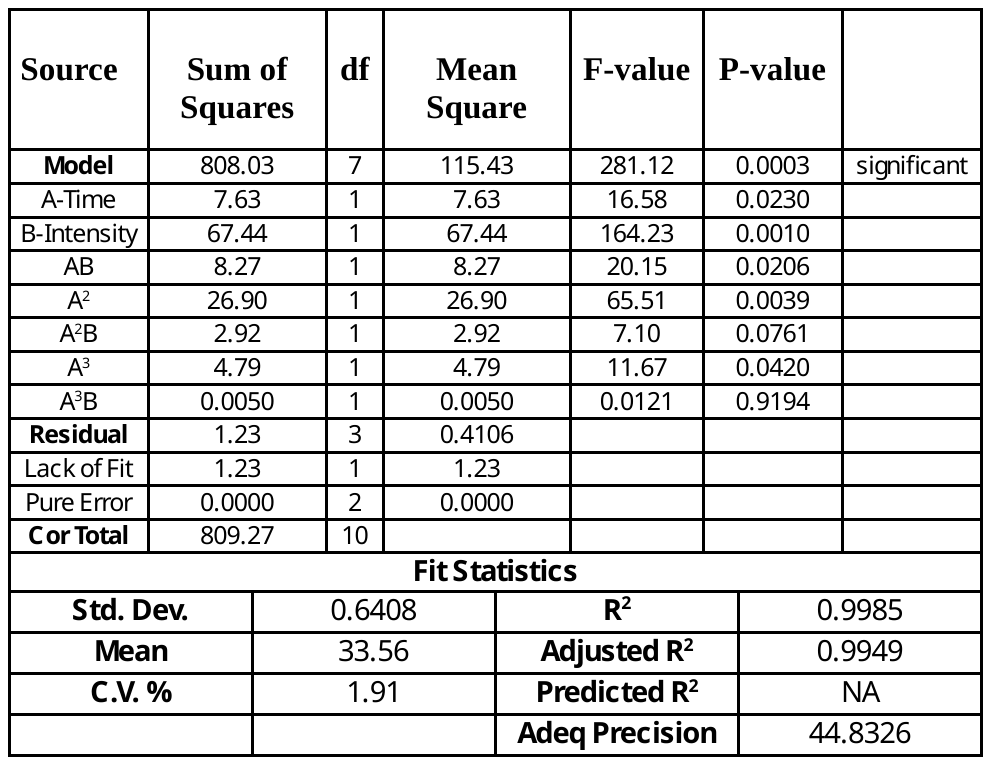

**Table S2.** Analysis of variance (ANOVA) for the reduced quartic model evaluating the effects of ultrasound time (A), intensity (B), and frequency (C) on total reactive oxygen species (ROS) generation during acellular sonodynamic therapy (SDT) experiments. The model shows significant predictive capability (P = 0.0316) with acceptable goodness-of-fit (R² = 0.9379, Adeq Precision = 10.39).

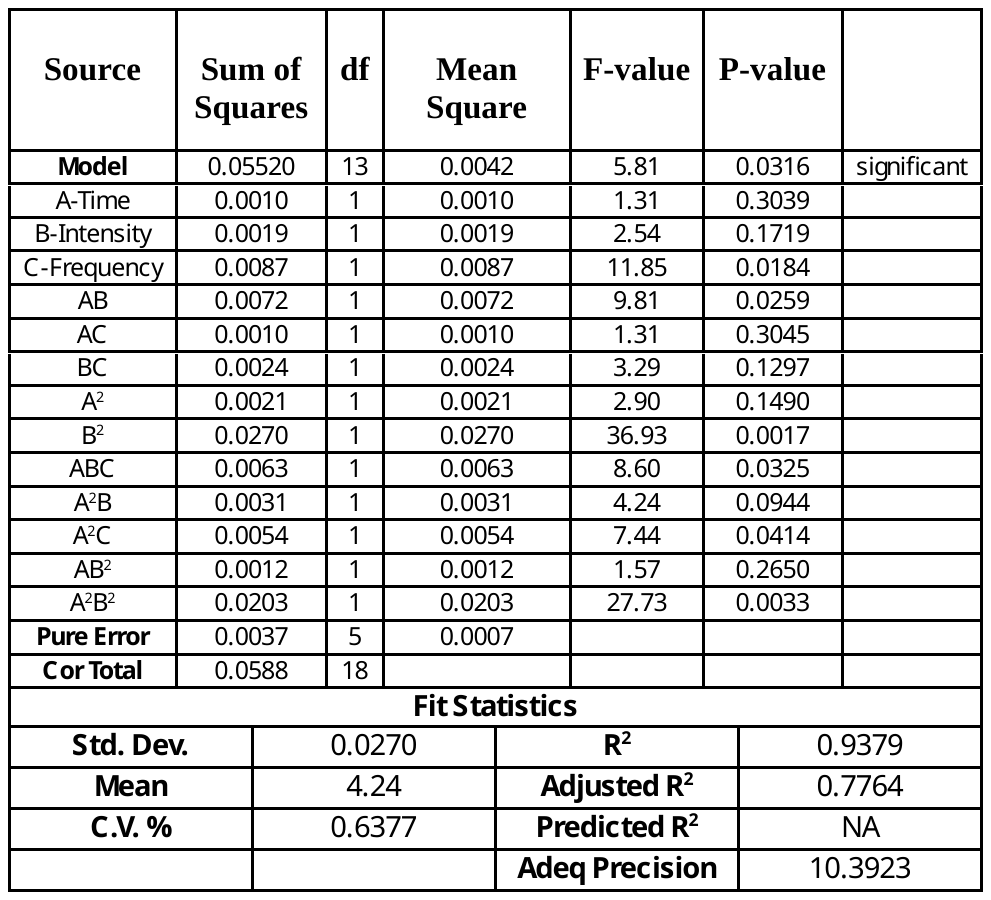
